# A modular circuit architecture coordinates the diversification of courtship strategies in *Drosophila*

**DOI:** 10.1101/2023.09.16.558080

**Authors:** Rory T. Coleman, Ianessa Morantte, Gabriel T. Koreman, Megan L. Cheng, Yun Ding, Vanessa Ruta

## Abstract

Identifying a mate is a central imperative for males of most species but poses the challenge of distinguishing a suitable partner from an array of potential male competitors or females of related species. Mate recognition systems are thus subject to strong selective pressures, driving the rapid coevolution of female sensory cues and male sensory preferences. Here we leverage the rapid evolution of female pheromones across the *Drosophila* genus to gain insight into how males coordinately adapt their detection and interpretation of these chemical cues to hone their mating strategies. While in some *Drosophila* species females produce unique pheromones that act to attract and arouse their conspecific males, the pheromones of most species are sexually monomorphic such that females possess no distinguishing chemosensory signatures that males can use for mate recognition. By comparing several close and distantly-related *Drosophila* species, we reveal that *D. yakuba* males have evolved the distinct ability to use a sexually-monomorphic pheromone, 7-tricosene (7-T), as an excitatory cue to promote courtship, a sensory innovation that enables *D. yakuba* males to court in the dark thereby expanding their reproductive opportunities. To gain insight into the neural adaptations that enable 7-T to act as an excitatory cue, we compared the functional properties of two key nodes within the pheromone circuits of *D. yakuba* and a subset of its closest relatives. We show that the instructive role of 7-T in *D. yakuba* arises from concurrent peripheral and central circuit changes: a distinct subpopulation of sensory neurons has acquired sensitivity to 7-T which in turn selectively signals to a distinct subset of P1 neurons in the central brain that trigger courtship behaviors. Such a modular circuit organization, in which different sensory inputs can independently couple to multiple parallel courtship control nodes, may facilitate the evolution of mate recognition systems by allowing males to take advantage of novel sensory modalities to become aroused. Together, our findings suggest how peripheral and central circuit adaptations can be flexibly linked to underlie the rapid evolution of mate recognition and courtship strategies across species.

## Main

Sensory evolution has been proposed to fuel behavioral diversification across species (*1*), allowing animals to capture and perceive distinct features of their environment. The rapid expansion and diversification of sensory receptors is thought to represent a potent force in the evolution of behavior, endowing each species with the ability to detect the signals relevant to their ecological niche or lifestyle (*1–5*). Changes to central circuit processing, however, also undoubtedly play a significant role in behavioral evolution (*6*, *7*), acting in concert with a diversifying periphery to integrate and interpret novel sensory inputs and mediate coherent behavioral responses. Yet, how peripheral and central circuit adaptations are coordinated to give rise to the emergence of novel behavioral traits remains unclear.

Reproductive behaviors provide a powerful inroad to explore how evolution acts at different levels within a sensory-processing circuit. As species diverge, the sensory signals females convey to males rapidly diversify to prevent interspecies mating (*8*). In turn, the sensory pathways that males use to both detect and interpret female cues must coevolve to underlie successful mate discrimination. Indeed, changes in mate preference between closely-related species are thought to often rely on reciprocal switches in the behavioral valence of mating signals (*9*)–conspecific cues must be made arousing while heterospecific cues made aversive–a process likely relying on concurrent changes in peripheral detection and central circuit processing.

The *Drosophila* genus offers an opportunity to examine how the rapid diversification of mate recognition systems unfolds. Male flies instinctively court females with an elaborate ritual that is highly stereotyped within a species (*10*), yet highly divergent across species (*11*, *12*). Many *Drosophila* species live in sympatry with a subset of their close relatives (*13*), driving the evolution of behavioral and morphological traits that signal species identity and contribute to mate recognition. In particular, cuticular pheromones have rapidly and repeatedly diversified across members of this genus, both in their chemical composition and the apparent logic of how they control mate choice (*14–19*) (**Fig. 1a**), suggesting that males must correspondingly have evolved distinct mechanisms to detect and interpret these chemical cues. In some species, females produce unique cuticular hydrocarbon compounds thought to act as sex pheromones that aid in mate recognition. For example, *D. melanogaster* females produce 7,11-heptacosadiene (7,11-HD), a pheromone that differs from those carried by conspecific males and heterospecific females that cohabitate within the same environments. 7,11-HD thus signals both the sex and species identity of *D. melanogaster* females, enabling it to serve as a potent cue to promote courtship by conspecific males. Sexually instructive pheromones like 7,11-HD, however, appear to be an exception in the *Drosophila* genus as females in the vast majority of species do not appear to produce any chemicals that uniquely distinguish them from their conspecific males (*15*)(**Fig. 1a**, 84 of 99 species sampled). For example, *D. yakuba* and *D. simulans,* close relatives believed to have evolved in geographic separation (*20*), have independently lost the biosynthetic enzymes necessary to produce female-specific diene pheromones like 7,11-HD (*19*, *21*, *22*). As a consequence, both *D. yakuba* and *D. simulans* females produce predominantly 7-tricosene (7-T) (**Fig. 1a**), the same cuticular compound as their males. *D. simulans* males appear insensitive to 7-T (*17*), likely relying on vision or other sensory cues to become aroused and pursue their conspecific females. Instead, pheromone processing pathways in *D. simulans* have been proposed to be largely dedicated to the detection of inhibitory pheromones, like 7,11-HD, to suppress futile courtship of heterospecific females (*17*). The prevalence of species with sexually-monomorphic pheromones throughout the *Drosophila* genus (**Fig. 1a**) suggests that *D. simulans* may represent a conserved sensory logic for mate recognition, in which males rely on vision rather than chemosensation to become aroused and only depend on pheromones as inhibitory signals to prevent heterospecific courtship. Alternatively, given the evolutionary flexibility of mate recognition systems, males of other monomorphic species may have evolved novel mechanisms to use sexually-ambiguous pheromones to become aroused and trigger their courtship.

**Figure 1.**
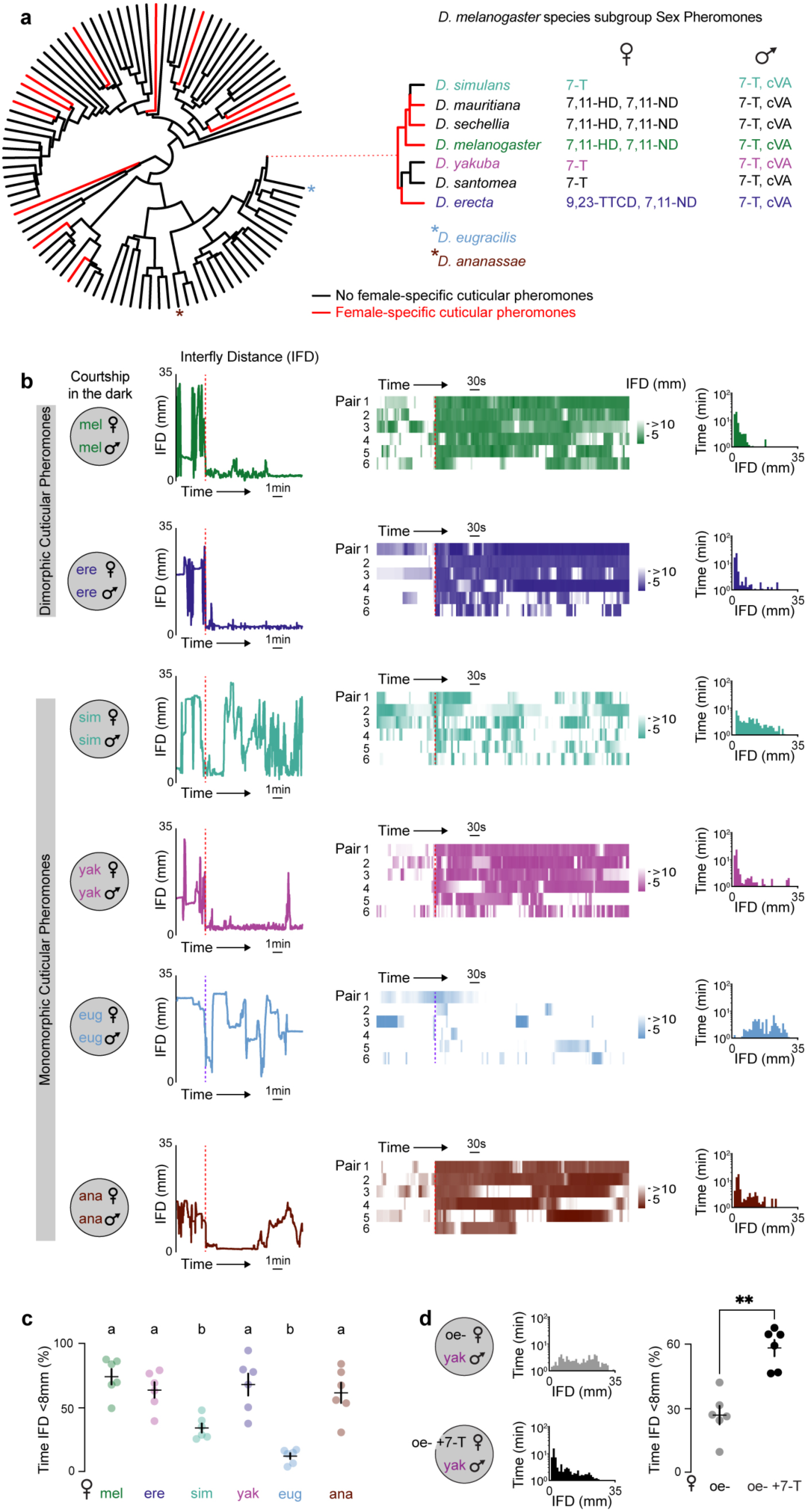
Sexually ambiguous pheromones do not preclude courtship in the dark. **a,** Phylogeny of 99 *Drosophila* species where cuticular pheromones have been characterized (left) and the primary sex pheromones of the species of the *D. melanogaster* subgroup (right) (*14*, *15*). 7-T: 7-tricosene; cVA: cis-Vaccenyl acetate; 7,11-HD: 7,11-heptacosadiene; 7,11-ND: 7,11-nonacosadiene; 9,23-TTCD: 9,23-tritriacosadiene. **b**, Courtship as captured by the inter-fly distance (IFD) between a *D. melanogaster* (mel), *D. erecta* (ere), *D. simulans* (sim), *D. yakuba* (yak), *D. eugracilis* (eug), or *D. ananassae* (ana) male with a conspecific female in the dark. (Left) IFD traces over time for a single representative pair as courtship proceeds. (Middle) Heatmaps for 6 pairs, aligned to courtship initiation for all species except *D. eugracilis* where initiation of courtship in the dark was never observed and purple dotted line marks the time of first interaction. (Right) Histograms of the time as a function of IFD for same 6 courting pairs. **c**, Average percent time pairs from **b** spent at IFD<8mm after first interaction (eug) or after courtship initiation (all other species). Each data point represents the individual pairs shown in the heatmaps in (b). **d**, Histograms and average percent time *D. yakuba* males spent at IFD<8mm with oenocyte-less females mock perfumed (oe-) or perfumed with the *D. yakuba* pheromone 7-T in the dark. Statistical tests performed were ANOVA with Tukey’s post-hoc (c) or unpaired Mann-Whitney (d). Data points represent individual males and error bars are mean±SEM. Letters above data sets denote statistically different groups (p<0.05). Asterisks denote **P<0.01.

Here we explore how sexually-monomorphic pheromones are used to guide mate recognition across species. We find that despite the similar evolutionary trajectories of *D. simulans* and *D. yakuba, D. yakuba* males have adopted a distinct strategy in which they use their sexually-ambiguous pheromone 7-T, as an excitatory cue that promotes courtship. This sensory innovation confers *D. yakuba* males with the ability to become aroused and faithfully pursue their conspecific females in the dark, thereby expanding the potential sensory environments available for mating. The distinct behavioral sensitivity of *D. yakuba* males to 7-T arises from coordinated peripheral and central circuit adaptations: 7-T activates a distinct subset of sensory neurons, which selectively signal to one of two molecularly-defined subsets of P1 neurons that control a male’s sexual arousal (*23–35*). While activation of either P1 neuron subtype in *D. yakuba* males is sufficient to evoke courtship behaviors, only the P1 subpopulation tuned to 7-T is necessary for *D. yakuba* males to maintain pursuit of females in the dark. *D. yakuba* males thus display striking differences from *D. simulans* in how they detect and integrate pheromone signals, allowing them to potentially occupy a distinct sensory niche for mating. A similar sensory specialization is also apparent in the P1 subpopulations of *D. melanogaster* males, underscoring how the modular organization of this circuit node may provide a facile evolutionary substrate for the rapid diversification of pheromone preferences and courtship strategies.

### *D. yakuba* males are aroused by a sexually-monomorphic pheromone

In many *Drosophila* species, the perception of a moving fly-sized target serves as a potent courtship-promoting cue, suggesting that vision can act redundantly with female pheromones to arouse males (*16*, *33*, *34*, *36*). To compare the role played by chemosensory signaling in mate recognition across species, we examined their courtship dynamics in the dark, where mate recognition becomes more reliant on chemical cues (*37*). We focused our comparison on two species that produce different sexually-dimorphic cuticular pheromones—*D. melanogaster* and *D. erecta*—and several in which females lack any chemical signatures that differentiate them from their males—*D. simulans,* a sister species of *D. melanogaster; D. yakuba*, another close relative within the *D. melanogaster* species subgroup; *D. eugracilis*, a close outgroup species, and *D. ananassae*, a more distant relative (**Fig. 1a**). Given that a male’s faithful pursuit of a female represents a conserved hallmark of male courtship throughout this genus (*10*, *11*), we used inter-fly distance (IFD) over time as a readout of courtship independent of any potential variation in motor displays across species (**Fig. 1b; Fig. S1a,b**).

*D. melanogaster* males have been shown to court proficiently in the dark (*37–41*), relying on 7,11-HD, the pheromone carried by their conspecific females to sustain their arousal in the absence of vision (*37*, *39*, *40*). Indeed, once *D. melanogaster* males initiated courtship, they closely tracked a target female, modulating their locomotion to keep her in close proximity (<8 mm) for bouts lasting many minutes (**Fig. 1b,c; Fig. S1b,d; Supp. Video 1**). *D. erecta* males displayed similarly robust pursuit of their conspecific females, consistent with the notion that the unique pheromonal cues on *D. erecta* females likewise signify the presence of an attractive mate and can promote courtship even in the absence of vision (**Fig. 1b,c**). In contrast, the fidelity of a male’s courtship pursuit was far more variable across monomorphic species. *D. eugracilis* males never appeared to perform any bouts of courtship in the dark, and proximity to the female was transient and incidental, suggesting a strict requirement for vision (**Fig. 1b,c**). *D. simulans* males displayed brief bouts of courtship, but these frequently broke off as soon as the female walked away (**Fig. 1b,c; Fig. S1a,b,d; Supplemental Video 2**), indicating males are unable to sustain their pursuit without visual feedback. In contrast, *D. yakuba* and *D. ananassae* males engaged in extended periods of courtship pursuit, resembling the persistent tracking displayed by species with sexually-dimorphic pheromones (**Fig. 1b,c; Fig. S1a,b,d; Supp. Video 3**). Thus it appears that the males of some monomorphic species, like *D. yakuba* and *D. ananassae,* may have evolved distinct sensory strategies to identify and pursue their conspecific females in the absence of visual feedback or a female-specific pheromone, underscoring how variation in sensory signals can guide mating decisions (*42*, *43*).

In *D. melanogaster* males, the excitatory effect of 7,11-HD is thought to compensate for the absence of vision (*37*, *39*, *40*). The persistent courtship pursuit exhibited by *D. yakuba* males in the dark suggests the possibility that they may similarly rely on female pheromones to become aroused, despite the sexually-monomorphic nature of 7-T. Indeed, in the dark, *D. yakuba* males were largely indifferent to *D. melanogaster* females that lack oenocytes (oenocyte-less [oe-]) and thus do not produce cuticular pheromones (*16*). However, perfuming oe-females with 7-T rendered them attractive and elicited extended bouts of courtship pursuit (**Fig. 1d; Fig. S1c,e**). The detection of 7-T therefore appears sufficient to arouse *D. yakuba* males, an adaptation that could expand the diurnal periods accessible for mating, consistent with their preference to court before dawn (*38*). Consistent with the arousing role of 7-T, *D. yakuba* males vigorously courted *D. simulans* females to the same extent as their conspecifics (**Fig. S1d**). 7-T thus appears to be used in a divergent manner to hone mate selection, serving as an inhibitory cue to prevent heterospecific courtship in *D. melanogaster* males (*44*), a neutral signal that *D. simulans* males appear indifferent to (*17*), and an excitatory cue that promotes courtship in *D. yakuba* males, reflecting multiple switches in the behavioral valence of this pheromone across closely-related species.

### P1 neuron pheromone tuning has rapidly diversified across species

P1 neurons serve as a central node within the male courtship circuit, integrating from multiple sensory pathways to encode the suitability of a prospective mate and trigger persistent courtship behaviors (*23–35*). The distinct reliance of different *Drosophila* species on pheromone signals to guide their courtship (**Fig. 1b**) suggests the possibility that the chemosensory tuning of P1 neurons has accordingly diversified to mediate species-specific mate preferences. We therefore generated neurogenetic tools in *D. simulans* (*17*)*, D. yakuba* (*45*), and *D. erecta* to allow a comparative analysis of the pheromone tuning of the P1 population. Anatomic labeling revealed that the P1 neurons in all four species displayed rich projections in the lateral protocerebral complex (LPC), a sexually-dimorphic neuropil that in *D. melanogaster* has been shown to receive input from multiple sensory-processing pathways and extend outputs to descending neurons that drive the component behaviors of male courtship (*25*, *26*, *46*)(**Fig. 2a**). Minor differences in the arborization patterns of P1 neurons were apparent across species. Yet despite their morphological variation, we found that P1 neurons nevertheless played a common role in promoting courtship across species as optogenetic activation of this population in *D. erecta* and *D. yakuba* males was sufficient to trigger robust and persistent pursuit of otherwise unattractive heterospecific female targets (**Fig. S2a,b**), replicating the courtship-promoting function of these neurons in *D. melanogaster* and *D. simulans* (*17*, *24*, *25*, *27*, *29*, *30*). P1 neurons thus represent a conserved circuit node regulating male courtship.

**Figure 2.**
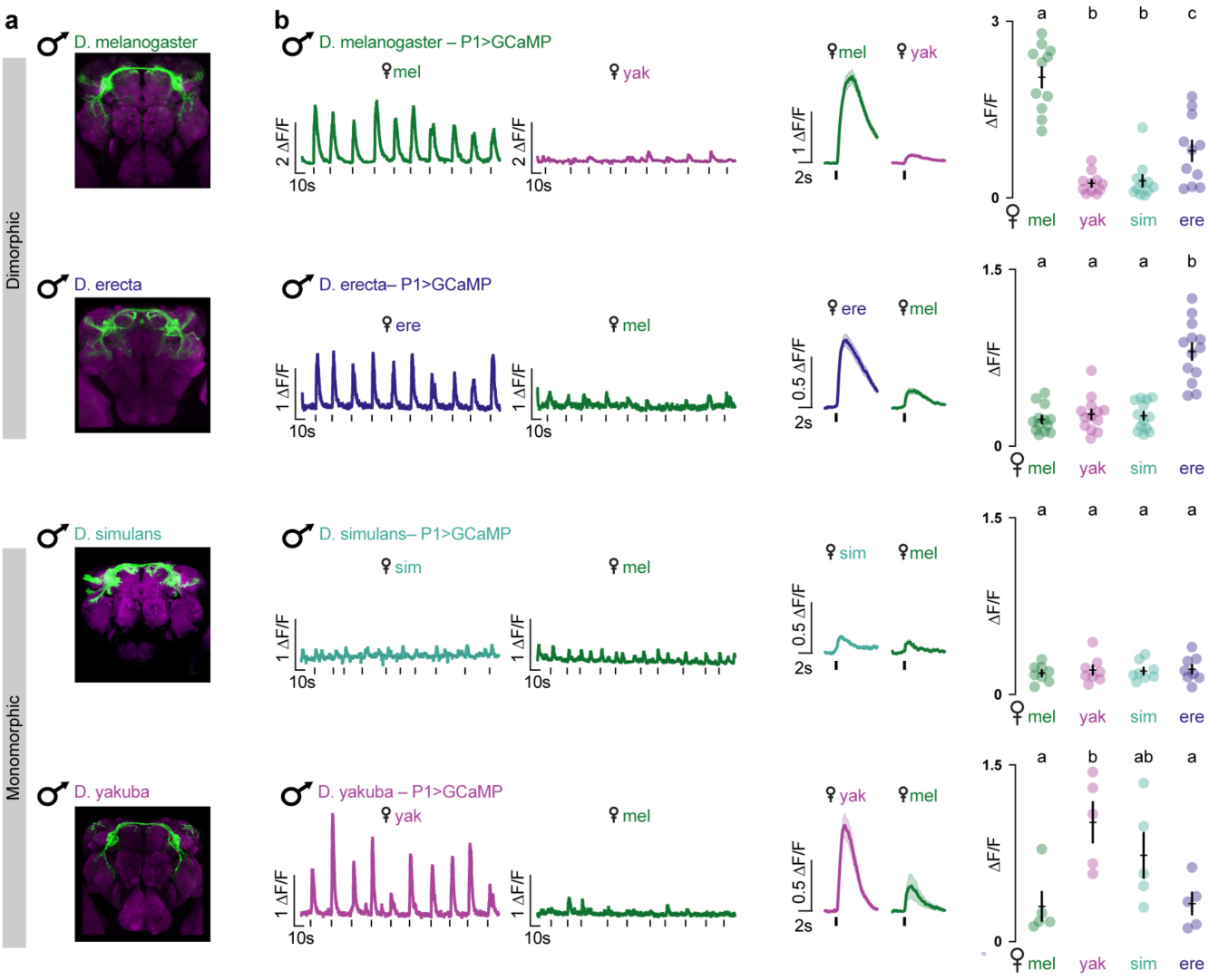
P1 neurons of *D. yakuba* males share conspecific tuning pattern of dimorphic species. **a**, P1 neurons labeled by 71G01>CD8::GFP (*D. melanogaster, D. erecta*), 71G01>GCaMP6s (*D. simulans*), or SplitP1>CD8::GFP (*D. yakuba*) expression, stained for GFP (green) and neuropil counterstain (magenta). Images were masked to remove glial fluorescence from ie1 marker and non-P1-specific labeling for clarity. **b**, (Left) Representative traces of P1 responses in the LPC in males of each species evoked in response to the taste of a conspecific or heterospecific female (black ticks indicate time of foreleg taps). (Middle) Averaged tap-evoked functional responses (ΔF/F_0_, black ticks) of the P1 neurons. (Right) Average peak response (ΔF/F_0_) for each male evoked by a given female target. Statistical tests performed were ANOVA with Tukey’s post-hoc. Data points represent individual males and error bars are mean±SEM. Letters above data sets denote statistically different groups (p<0.05).

To explore how the pheromonal responses of P1 neurons vary across species, we performed functional calcium imaging of P1 neuron projections within the LPC of tethered males, as they walked on an air-supported ball and were offered an array of female targets, replicating the sensory sampling males perform as they encounter females in the dark (**Fig. 2b**). In *D. melanogaster* (*17*, *25*, *28*) and *D. erecta* males, P1 neurons were robustly activated each time a male tapped his conspecific female, while minimal excitation was elicited by the taste of heterospecific females (**Fig. 2b**). This selective chemosensory tuning is consistent with the notion that the unique pheromones carried by females in sexually-dimorphic species serve as instructive cues for mate recognition, exciting the P1 neurons to promote courtship. In contrast, the P1 neurons of *D. simulans* males were unresponsive to the taste of any female targets (**Fig. 2b**), concordant with evidence that in this species, pheromone pathways do not appear to drive courtship of conspecific females and instead suppress inappropriate pursuit of heterospecific females (*16*, *17*). The P1 neurons of *D. yakuba* males, however, rather than replicating the attenuated chemosensory responses of *D. simulans,* instead resembled the selective tuning of species with sexually-dimorphic cuticular pheromones: the P1 neurons of *D. yakuba* males were strongly activated by the taste of both *D. yakuba* and *D. simulans* females but only weakly responsive to heterospecific female targets (**Fig. 2b**), consistent with behavioral evidence that 7-T serves as an excitatory cue sufficient to arouse *D. yakuba* males (**Fig. 1d**). Thus, while P1 neurons play a conserved role in promoting courtship across species, their pheromonal tuning has diversified, highlighting the intrinsic evolutionarily flexibility of sensory circuits that control mate recognition.

### Diversification of peripheral pheromone responses in *D. yakuba*

Together, our functional and behavioral data suggest that *D. yakuba* males have evolved a mechanism to use the sexually ambiguous chemical 7-T as an instructive courtship cue via the diversification of pheromone circuits that ultimately impinge onto P1 neurons. The distinct chemosensory strategy of *D. yakuba* males could reflect differences in the way 7-T is detected at the sensory periphery, variation in how this pheromonal signal is integrated by the P1 neurons, or both. To differentiate between these possibilities, we examined the peripheral sensory populations mediating pheromone detection in *D. yakuba,* to gain insight into how their diversification may contribute to 7-T acquiring an excitatory role in this species.

Cuticular pheromones in *Drosophila* are thought to be detected by a heterogenous sensory population marked by expression of the DEG/ENaC channel, Ppk23, which serves as an essential component of the pheromone transduction machinery (*17*, *40*, *47–50*). In *D. melanogaster*, Ppk23+ neurons adopt a paired organization within the sensory bristles of the male foreleg (*26*, *39*, *40*, *47*, *48*, *51*), in which one neuron co-expresses the DEG/ENaC channel Ppk25 and is tuned to 7,11-HD to promote courtship of conspecific females, while its Ppk25-partner is responsive to heterospecific pheromones, including 7-T, to suppress interspecies courtship (**Fig. 3a,h**). To explore the role that Ppk23-mediated pheromonal signaling plays in mate recognition in *D. yakuba,* we used CRISPR/Cas9 genome editing to delete the gene encoding this channel and assessed the courtship of mutant males paired with either conspecific or heterospecific females (**Fig. S3a)**. We found that *D. yakuba* males mutant for *ppk23* lost their characteristic aversion to heterospecific female targets, supporting that, as in *D. simulans* (*17*) this channel mediates the detection of inhibitory pheromones to curb inappropriate visual pursuit (**Fig. 3b,h**). Although *D. yakuba ppk23* mutant males vigorously pursued their conspecific females in the light, they were unable to sustain courtship in the dark, and their pursuit instead mirrored the saltatory dynamics displayed by *D. melanogaster ppk23* mutant males in the absence of visual feedback (**Fig. 3c,d; Fig. S3b,c**). Ppk23 signaling in *D. yakuba* thus appears to be required both to promote courtship of conspecific females and to inhibit pursuit of heterospecific targets suggesting that, as in *D. melanogaster* (*40*, *47*, *48*), Ppk23 receptors mark a heterogeneous neuronal population that plays opposing roles in honing mate recognition.

**Figure 3.**
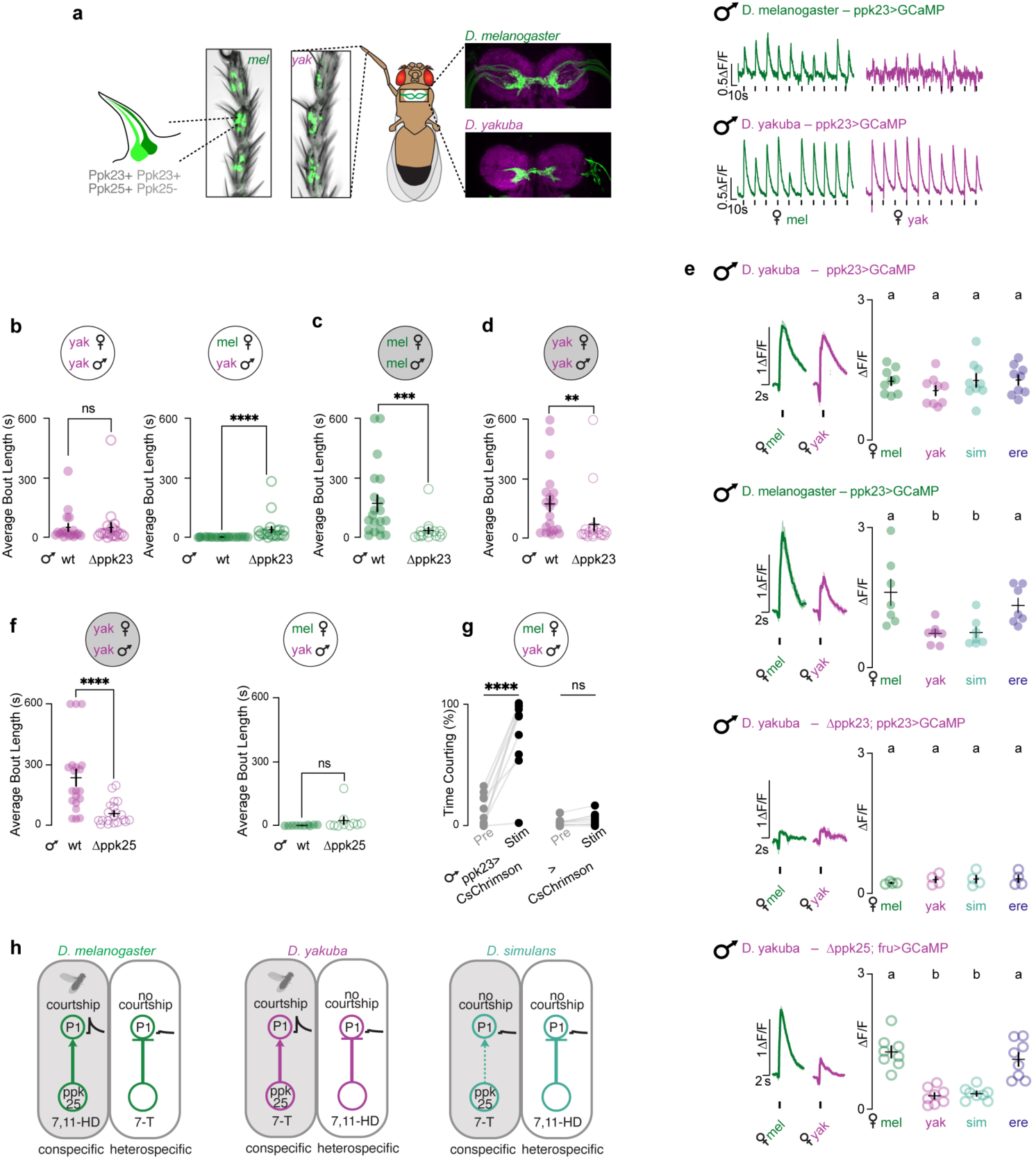
Altered pheromone sensitivity in *D. yakuba* sensory neurons. **a,** (Left) Cartoon showing the organization of paired Ppk23+ sensory neurons within chemosensory bristles and bright field image showing expression of CD8::GFP (green) in the soma of the Ppk23+ sensory neurons in the foreleg tarsal segments of *D. melanogaster* and *D. yakuba* males. Images are distal end up. (Middle) Cartoon showing the Ppk23+ sensory afferents in the ventral nerve cord (VNC) targeted for recording pheromone responses and anatomic images of Ppk23+ sensory afferents expressing CD8::GFP (green) with neuropil counterstain (magenta). Images are anterior side up. (Right) Representative functional responses of Ppk23+ sensory afferent in a *D. melanogaster* or *D. yakuba* male evoked by the taste of a *D. melanogaster* (green) or *D. yakuba* (purple) female (black tick marks indicate time of foreleg taps). **b,** Average courtship bout length in the 10 minutes following courtship initiation for wildtype or *ppk23* mutant *D. yakuba* males towards *D. yakuba* or *D. melanogaster* females in the light. **c,d**, Average courtship bout length of *ppk23* mutant *D. melanogaster* (c) or *D. yakuba* (d) males towards a conspecific females in the dark. **e,** (Left) Average functional responses (ΔF/F_0_) aligned to time of a tap of indicated conspecific or heterospecific female of the foreleg sensory afferents and (Right) average peak response (ΔF/F_0_) per male for a given female target. **f**, Average courtship bout length of wildtype and *ppk25* mutant males toward *D. yakuba* females in the dark (left) and *D. melanogaster* females in the light (right). **g**, Percent time courting prior to (Pre) or during (Stim) periods of optogenetic stimulation of *D. yakuba* males expressing CsChrimson in Ppk23+ sensory neurons (ppk23>CsChrimson) or control animals (>CsChrimson) paired with a *D. melanogaster* female. **h,** Proposed model summarizing the inferred or observed changes in the sensitivity of peripheral sensory populations and the resulting effect on P1 neuron activity. Dotted line represents signaling inferred from previously reported behavioral data (*52*). Shading in response traces represents mean±SEM. Data points represent individual males and error bars are mean±SEM. Statistical tests performed were ANOVA with Tukey’s post-hoc (e), unpaired Mann-Whitney (b-d, f), or paired t-test (g). Letters above data sets denote statistically different groups (p<0.05). Asterisks denote **P<0.01, ***P<0.001, ****P<0.0001, ns, not significant.

To gain insight into how the chemical tuning of the Ppk23+ sensory population relates to their dual role in promoting and suppressing courtship, we generated a *ppk23-Gal4* driver in *D. yakuba* to gain genetic access to the full sensory ensemble (**Fig. 3a**). While the forelegs of *D. melanogaster* and *D. yakuba* males harbored comparable numbers of Ppk23+ neurons that maintained a similar paired arrangement (**Fig. S3d**), they displayed striking differences in their pheromone sensitivities. Imaging the aggregate activity of Ppk23+ foreleg sensory afferents in *D. yakuba* males revealed that they were robustly activated by all target flies, with equivalent responses evoked by the taste of both conspecific and heterospecific females (**Fig. 3e**). Such broad tuning contrasts with the selective responses to 7,11-HD-carrying females observed in both *D. melanogaster* (**Fig. 3e**) and *D. simulans* males (*17*), to alternatively promote or deter courtship to *D. melanogaster* females (*17*) (**Fig. 3h).** We confirmed that *D. yakuba* Ppk23+ sensory responses were chemosensory in origin, as oe-females lacking pheromones evoked significantly weaker activity but robust responses could be restored by perfuming these females with either 7,11-HD or 7-T (**Fig. S3e**). Moreover, pheromone responses in *D. yakuba* males were largely abolished in *ppk23* mutants (**Fig. 3e**), substantiating the conserved and essential role that Ppk23- mediated signaling plays in pheromone detection across species (*17*, *40*, *47*, *48*).

To determine whether the broad chemosensory tuning of Ppk23+ neurons reflects the activity of heterogenous Ppk25+ and Ppk25- subpopulations (**Fig. 3a**), we generated a *ppk25* mutant (**Fig. S4a**). Recording from foreleg sensory afferents in *ppk25* mutant males revealed that responses to *D. yakuba* and *D. simulans* females were selectively attenuated, while responses evoked by other heterospecific female targets remained intact (**Fig. 3e**), suggesting that a Ppk25- mediated pathway underlies 7-T detection to regulate a male’s sexual arousal. Consistent with this proposal, *D. yakuba ppk25* mutants displayed diminished courtship towards conspecifics in the dark (**Fig. 3f; Fig. S4b**), phenocopying the disrupted pursuit observed in *ppk23* mutants (**Fig. 3d**). *D. yakuba ppk25* mutants nevertheless remained averse to courting *D. melanogaster* females (**Fig. 3f**), substantiating that *ppk25* is required for detection of 7-T without impairing the recognition of heterospecific pheromones like 7,11-HD that suppress courtship.

Together, these observations suggest that the sensory neurons that detect conspecific and heterospecific pheromones have undergone a reciprocal switch in their chemical specificity across species such that in both *D. melanogaster* and *D. yakuba* males the Ppk25+ subpopulation mediates the detection of the distinct pheromones carried by their conspecific females to promote courtship (*26*, *49*), while the Ppk25- subpopulation underlies the detection of heterospecific pheromones. This swap in the chemical tuning of peripheral sensory neuron subtypes suggests a potentially facile mechanism to alter the behavioral meaning of pheromones, whereby the ascending circuits that promote or suppress courtship are conserved but their pheromone sensitivity is altered. In *D. simulans* males*, ppk25* has been reported to promote courtship (*52*), suggesting it may be similarly involved in 7-T detection. However, this peripheral sensitivity does not appear to be translated to P1 neuron excitation (**Fig. 2b; 3h**), in accord with the tepid courtship that males of this species display in the dark (**Fig. 1b,c**). The behavioral valence of pheromones may therefore depend, not only on which subset of sensory neurons are activated, but also how these peripheral signals are conveyed to the P1 population. Indeed, using optogenetics to exogenously activate the Ppk23+ sensory neurons and bypass pheromone detection revealed that while stimulation of this population suppresses courtship in *D. simulans* (*17*), it drove *D. yakuba* males to indiscriminately court heterospecific females (**Fig. 3g**), replicating their courtship-promoting role in *D. melanogaster* males (*17*, *40*, *47*). The opposing behavioral valence of Ppk23+ sensory neuron activation across species implies that, beyond the diversification of their pheromone tuning, additional neural adaptations underlie how these peripheral signals are transmitted to the P1 neurons to control a male’s mating decisions.

### Sensory specialization of molecularly distinct P1 populations

How might P1 neurons accommodate the flexible integration of different ascending pheromone pathways to underlie their species-specific tuning? The modular organization of sensory circuits has been proposed to facilitate their evolutionary diversification, as their segregated nature allows for the independent retuning of sensory inputs with minimal pleiotropic effects (*53*, *54*). Notably, P1 neurons can be divided into two discrete populations that are differentially marked by the expression the sexually-dimorphic transcription factors, *doublesex* (*dsx*) and *fruitless (fru)*. Both populations play a concerted role in regulating courtship (*23*, *55*– *57*), suggesting that these distinct P1 subsets may represent modular units that could integrate from different sensory channels and confer additional flexibility to how ascending pheromone signals are weighted.

To explore this possibility, we used intersectional genetic strategy to label the Fru+ (Fru∩P1) or Dsx+ (Dsx∩P1) P1 populations (**Fig. S5a**), each of which marks a comparable number of neurons in *D. yakuba* and *D. melanogaster* males (**Fig. S5b-e**). While the morphology of the Fru∩P1 and Dsx∩P1 subpopulations was largely similar both within and across species, minor variation was apparent in their ventral and dorsal lateral projections (**Fig. 4a,b; Fig. S5b,c,f,g**), suggesting they are anatomically poised to receive distinct inputs. Consistent with this, we observed striking differences in the functional tuning of these P1 neuron subsets in *D. yakuba* males. Dsx∩P1 neurons were strongly activated by the taste of conspecific females (**Fig. 4b**), which we confirmed using perfumed oe-females reflects their distinct chemosensory tuning to 7-T (**Fig. S6a**). Fru∩P1 neurons, by contrast, were unresponsive to the taste of all target flies (**Fig. 4a**). Optogenetic activation of either neuronal population was sufficient to trigger courtship towards heterospecific females (**Fig. 4a,b**), although we observed minor quantitative differences in which component behaviors were evoked (**Fig. 4a,b; Fig. S7a,b**). These findings indicate that genetically distinct and functionally specialized subsets of P1 neurons play overlapping roles in triggering sexual arousal to drive male courtship, even towards inappropriate targets.

**Figure 4.**
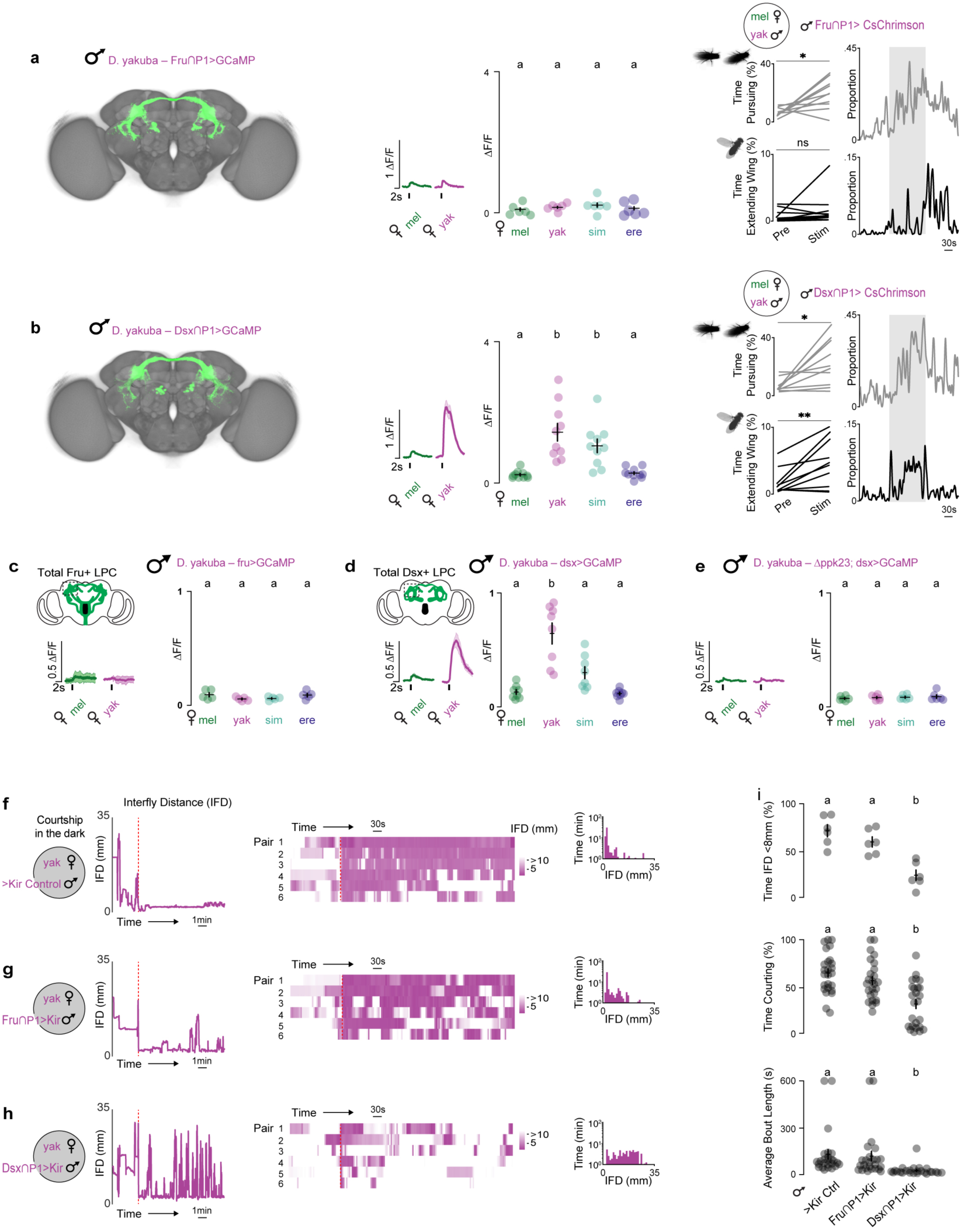
Sensory specialization of Fru+ and Dsx+ P1 subpopulations in *D. yakuba*. **a,b,** (Left) Maximum intensity projections of neurons labeled by intersection of Fru (a) or Dsx (b) and the P1-driver 71G01 (Fru∩P1 and Dsx∩P1, respectively) in *D. yakuba*, registered to a common template brain (see methods). (Middle) Averaged functional responses (ΔF/F_0_) aligned to time of a tap of indicated conspecific or heterospecific female and average peak response (ΔF/F_0_) per male for a given female target. (Right) Total percent time pursuing (top) or performing unilateral wing extensions (bottom) toward a *D. melanogaster* female target prior to (Pre) or during (Stim) optogenetic stimulation of Fru∩P1>CsChrimson or Dsx∩P1>CsChrimson males and the proportion of flies engaging in these behaviors over the course of the experiment. **c,d**, Functional responses of the Fru+ (c) or Dsx+ (d) neurons innervating the LPC where P1 neurons reside. (Left) Averaged functional responses (ΔF/F_0_) aligned to time of a tap of indicated conspecific or heterospecific female and (Right) average peak response (ΔF/F_0_) per male for a given female target. **e,** Functional responses of Dsx+ neurons in the LPC (as in d) in *ppk23* mutant males. **f,g,h**, Courtship in the dark as captured by inter-fly distance (IFD) of a *D. yakuba* male with constitutively silenced P1 subsets (Fru∩P1>Kir (g), Dsx∩P1>Kir (h), or genetic control (f; 71G01-DBD; UAS-Kir))towards a conspecific female in the dark. (Left) IFD traces over time for a single representative pair as courtship proceeds. (Middle) Heatmaps for 6 pairs, aligned to courtship initiation (red dotted line). (Right) Histograms of the time as a function of IFD for same 6 courting pairs. **i**, Percent time males spent at IFD<8mm after courtship initiation for 6 pairs in f-h, and percent time courting for 10 minutes after first initiation and average bout length for a larger (n=25) manually scored sample of each genotype. Shading in response traces represents mean±SEM. Data points represent individual males and error bars are mean±SEM. Statistical tests performed were ANOVA with Tukey’s post-hoc (**a-e, i**) or paired t-test (**a,b**). Letters above data sets denote statistically different groups (p<0.05). Asterisks denote *P<0.05, **P<0.01, ns, not significant.

Given that all P1 neurons are thought to express Dsx, with only a subset co-expressing Fru (*23*, *55–57*)(**Fig. S5b-e**), the differential sensory tuning we observed suggests that the Dsx+/Fru-P1 population is selectively excited by 7-T in *D. yakuba* males. To capture the activity of the full population of Fru+ or Dsx+ P1 neurons independent of our intersectional labeling strategy, we used genetic drivers knocked into either the endogenous *fru* (*58*) or *dsx* locus and monitored pheromone responses in the lateral protocerebral complex (LPC) where P1 neuron processes reside. The aggregate activity all Fru+ or Dsx+ neurons replicated the pheromone tuning observed within each intersected P1 neuron subpopulation (**Fig. 4c,d**), with robust activation to the taste of *D. yakuba* females apparent only in the Dsx+ but not the Fru+ processes in the LPC. Responses to *D. yakuba* females were lost in *ppk23* mutants, supporting that the Dsx+ subpopulation uniquely integrates from the Ppk25+/Ppk23+ sensory pathways to underlie their excitation by 7-T (**Fig. 4e**). Dsx+ P1 neurons thus appear poised to play a distinct role in transducing pheromone detection to male arousal in *D. yakuba*. Consistent with this, constitutive silencing of the Dsx∩P1 neuronal subset strongly attenuated male courtship in the dark (**Fig. 4f,h,i**), replicating the abortive pursuit of *ppk23* and *ppk25* mutants (**Fig. 3d,f**). In contrast, silencing of the Fru∩P1 neuronal subset had little impact on courtship dynamics (**Fig. 4f,g,i**), underscoring the distinct role that these P1 subpopulations play in regulating courtship in the dark, where males become reliant on pheromonal feedback.

The molecular subdivision of P1 neurons by differential expression of the Fru and Dsx transcription factors represents a conserved feature of this circuit node (**Fig. S5b-e**), pointing to the possibility that this modular organization may serve as a more general substrate for the evolution of mate recognition in *Drosophila*. Indeed, in *D. melanogaster* males while activation of either Fru∩P1 and Dsx∩P1 neurons was sufficient to promote courtship (**Fig. S6b,c; S7c,d**), these P1 subsets displayed differential chemosensory tuning. Both P1 subpopulations responded robustly to their conspecific female pheromone (**Fig. S6b,c**), but the Fru+ subset was also strongly excited by the taste of *D. erecta* females (**Fig. S6b**), an expansion of pheromone tuning that may reflect sensitivity to 7,11-nonacosadiene (7,11-ND) a minor pheromone shared between *D. melanogaster* and *D. erecta* females (*14*)(**Fig. 1a**). The functional specialization of molecularly-defined P1 subtypes thus appears to be a conserved feature of courtship circuits across species, potentially facilitating the retuning and integration of new sensory pathways to allow for the emergence of species-specific mate preferences.

## Discussion

In this study, we leveraged the rapid evolution of female pheromones across the *Drosophila* genus to explore how changes in male pheromone detection and preference are coordinated to generate divergent mating strategies. Our work points to at least two discrete sites of diversification within the male courtship circuitry that can modify the behavioral valence of pheromones to underlie species-specific mate recognition. First, there has been a reciprocal swap in the chemical tuning of the Ppk25+ and Ppk25-sensory neurons within the male foreleg that detect conspecific and heterospecific pheromones and second, these peripheral signals are differentially conveyed to distinct subpopulations of P1 neurons, each sufficient to promote courtship (**Fig. 5a**). Such a modular circuit organization may facilitate the evolution of mate recognition systems by allowing rapidly diversifying peripheral sensory populations to independently couple to either of the P1 courtship control nodes, enabling males to take advantage of novel sensory modalities to drive courtship without compromising existing pathways for arousal (**Fig. 5b**). Exemplifying this evolutionary flexibility, we show that in *D. yakuba* males, one subset of P1 neurons uniquely integrates from the Ppk25+ peripheral sensory neurons with enhanced sensitivity to 7-T. The coordination of peripheral and central circuit modifications appears to allow *D. yakuba* males to use 7-T, a sexually-monomorphic and ambiguous compound, as an aphrodisiac that promotes courtship of conspecific females.

**Fig. 5.**
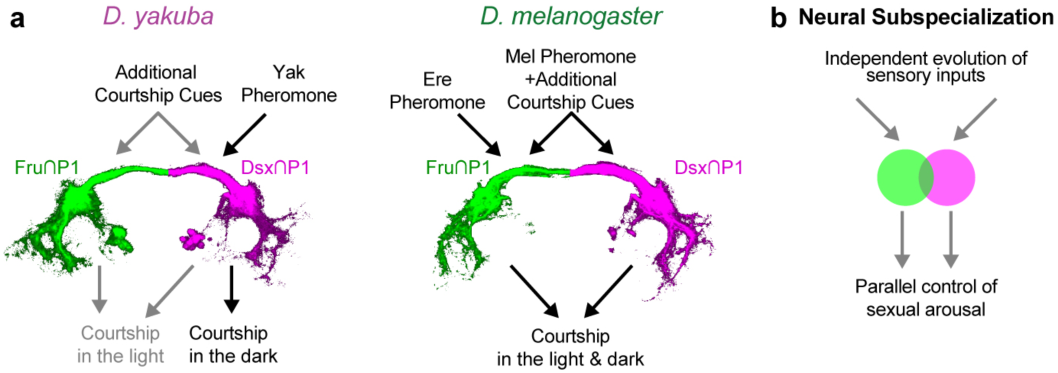
Subspecialization of P1 neuron subtypes. **a,** Diagram summarizing proposed sensory specializations of Fru∩P1 and Dsx∩P1 in *D. yakuba* and *D. melanogaster*. Pheromone inputs inferred from functional imaging and behavioral data in Fig. 4 & Fig. S6 are indicated by black arrows. Additional proposed sensory inputs and behavioral outputs are indicated by gray arrows. *D. melanogaster* (Mel), *D. yakuba* (Yak), *D. erecta* (Ere). **b**, Model for the diversification of courtship behaviors by neural subspecialization. Independent retuning of sensory inputs to behaviorally redundant but molecularly distinct P1 subtypes may facilitate the rapid evolution of sensory signals that control a male’s sexual arousal and courtship.

Female-specific pheromones would seem to offer obvious advantages to mate recognition, as they can serve as instructive cues that assure males become selectively aroused towards individuals of the appropriate sex and species (*59*). Yet, sexually-monomorphic pheromones are in fact representative of the vast majority of species within the *Drosophila* genus (*15*). For many *Drosophila* species, the visual perception of moving, fly-sized targets appears innately arousing, potentially rendering excitatory pheromones unnecessary to promote courtship (*16*, *33*, *34*, *36*). Nevertheless, as we reveal for *D. yakuba*, excitatory pheromones may confer additional robustness to courtship and expand the sensory environments in which it can occur. Indeed, by using 7-T to become aroused, *D. yakuba* males appear to overcome the strict requirement for vision, providing them with additional diurnal periods for mating (*38*). Notably, of the four monomorphic species we tested, *D. yakuba* and *D. ananassae* both robustly court in darkness, a sensory capacity shared by a number of distantly-related species distributed throughout this genus (*42*, *43*, *60–64*). The use of sexually monomorphic pheromones as generic courtship-promoting cues therefore appears to have arisen repeatedly, underscoring how the sensory pathways controlling courtship, likewise, have likely recurrently diversified to give rise to species with differential dependence on vision (**Fig. 1b, Fig. S8**).

While closely-related members of the *Drosophila* genus like *D. simulans* and *D. yakuba* have independently converged on the use of the same sexually-monomorphic pheromone profiles, our comparative analyses suggest that males of these species rely on distinct strategies for mate discrimination. The pheromone circuits of *D. simulans* males appear largely dedicated to the detection of heterospecific cues that suppress a male’s visual pursuit, thereby honing their courtship only towards appropriate females (*17*). Indeed, the *D. simulans* Ppk23+ sensory population shares the same selective tuning for 7,11-HD observed in *D. melanogaster* males, yet their activation suppresses courtship pursuit (*17*). In contrast, we find that the Ppk23+ population in *D. yakuba* males displays broad pheromone tuning, with robust responses to both conspecific and heterospecific pheromones, reflecting the dual use of pheromones as both excitatory and inhibitory cues in this species. Our data suggest that, as in *D. melanogaster* (*26*, *39*, *47*, *51*) expression of Ppk25 demarcates these opponent sensory populations (*26*, *39*, *47*, *51*), with Ppk25+ neurons sensitive to conspecific pheromones and the Ppk25-population responsive to heterospecific cues. The Ppk25+ population in *D. melanogaster* and *D. yakuba* have thus evolved distinct chemical sensitivities due to either the re-tuning of the receptors that detect pheromone ligands or alterations in their pattern of expression in the male foreleg. Ppk25 is not thought to be a component of the pheromone receptor complex that binds pheromonal compounds, suggesting it may instead mark and amplify excitatory pheromone pathways that promote courtship in different species (*50*, *51*). Indeed, the tepid courtship that *D. simulans* males display in the dark has been reported to be further attenuated in *ppk25* mutants (*52*). Our results thus underscore how the rapid evolution of female pheromones across the *Drosophila* genus may be partially accommodated by the evolution of peripheral sensory detection mechanisms, which must then be integrated with central circuits to drive mate recognition. Indeed in parallel to these peripheral adaptations, we that find that distinct subpopulations of P1 neurons differentially integrate from ascending pheromone pathways to contribute to the chemosensory control of courtship. While the structural or functional changes underlying the divergent patterns of P1 integration remain to be elucidated, notably the same ascending pathways that transmit pheromone signals from the foreleg afferents to the P1 neurons remain anatomically identifiable across species (*17*, *25*)(**Fig. S9**). Variation in the pheromone tuning of P1 neurons may therefore potentially arise from relatively minor adjustments to an otherwise conserved circuit architecture. Together, these data support the possibility that P1 neurons serves as a site of repeated evolutionary tinkering, whereby reweighting of the sensory input pathways to each P1 subpopulation would allow for the rapid emergence of species-specific mating strategies. The modular organization of molecularly distinct, behaviorally redundant P1 subtypes may further support their sensory diversification by ensuring that courtship is evolutionarily resilient, as one P1 subset can maintain ancestral patterns of mate regulation while novel sensory pathways are accommodated.

A key distinction between the functionally-specialized P1 neuron subsets is their differential expression of Fru and Dsx. As master regulatory transcription factors, Fru and Dsx act to shape the transcriptional landscape of the neurons in which they are expressed to define the sexually dimorphic features of the nervous system that confer the potential for sex-specific behaviors (*65–71*). Fru and Dsx act in a cell-autonomous manner to underlie the differences in neuron number, anatomy, and connectivity apparent across male and female brains (*24*, *72–75*), underscoring how the gain or loss of Fru or Dsx expression is poised to restructure the functional architecture of circuits. Notably, in contrast to the striking sexual dimorphisms in Fru+ and Dsx+ circuitry apparent within a species, these pathways appear largely conserved across males of different species (*17*, *76*, *77*), including in the numbers of neurons and arborization patterns of the different P1 subtypes. Thus, rather than changing patterns of Fru or Dsx expression, evolution may instead tinker with the anatomic or functional properties of neurons by altering the diverse gene expression programs that Fru and Dsx direct. The gain or loss of Fru or Dsx binding sites in downstream target genes offers a plausible and potentially powerful mechanism by which P1 subtypes could evolve species-specific sensory processing (*78–80*).

Our data suggest how behavioral evolution may proceed by neural subspecialization (**Fig. 5b**), a model with intriguing conceptual similarities to the mechanisms by which genes evolve through duplication and diversification. The functional redundancy of recently duplicated genes can facilitate the genetic drift of one or both copies, allowing for their subfunctionalization or neofunctionalization (*81*, *82*). Cellular duplication is not essential for neural subspecialization to arise. Rather, all that is required is a population of neurons with shared behavioral functions but with divergent molecular identities to allow neurons to independently diverge in their functional properties. Distinct molecular programs that specify the cellular properties of these neurons are necessary in order to provide the genetic substrate on which evolution can act. For example, the *cis*-regulatory landscape created by gain or loss of Fru-binding sites in P1 neurons may reshape the transcriptional programs controlling the connectivity or functional properties of the Fru+ subset, allowing for their independent diversification.

Our observations in *Drosophila* strengthen emerging evidence that modular neural circuit architectures, possibly arising from duplication and divergence, may represent a key evolutionary substrate for the diversification of complex behaviors (*54*, *83–88*). For example, the elaboration of cerebellar nuclei in different vertebrate species is thought to have arisen via repeated duplication of subnuclei and the transcriptional diversification of excitatory neurons to give rise to novel patterns of synaptic connectivity with their downstream targets (*85*). Similarly, the pallial-striatal circuits of songbirds mediating vocal learning have been proposed to evolve via the duplication and diversification of ancestral motor learning circuits (*83*, *84*). Central to these models of how behavioral circuits evolve is that duplication favors modular functional units which, as in gene duplication, relieves evolutionary constraints and allows for the diversification of the duplicated circuit structures. Here, by taking advantage of the rapid coevolution of the production of female pheromones and the pheromone preferences of males in a model clade (*6*, *89*), we have begun to map discrete sites of functional divergence to molecularly defined neural subpopulations, revealing neuronal modularity as an important substrate for behavioral evolution.

---

**Figure S1.**
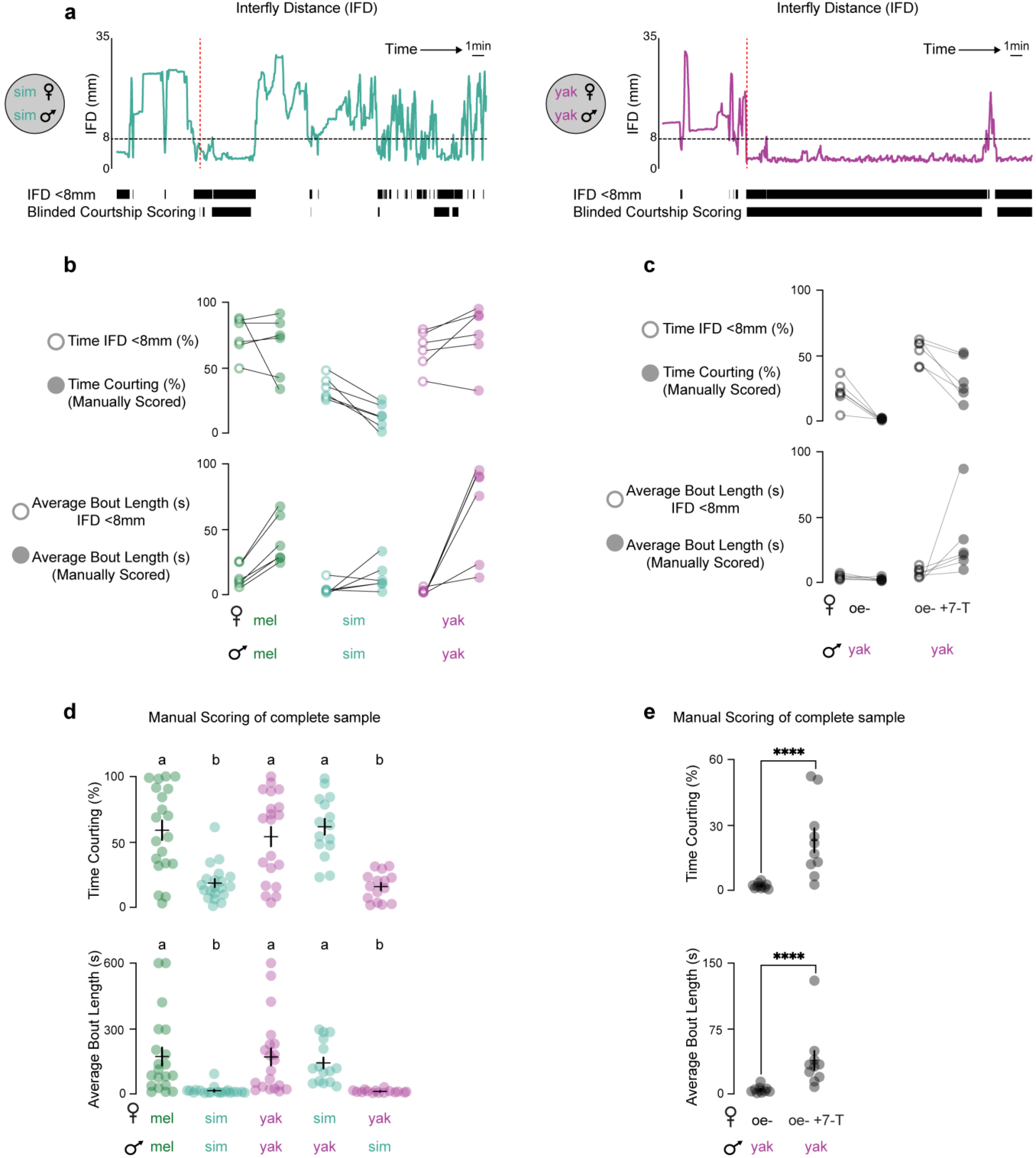
Inter-fly distances below 8mm approximate courtship in the dark. **a**, (Top) Inter-fly distance (IFD) traces over time for a single representative pair of *D. simulans* or *D. yakuba* as courtship proceeds. (Bottom) Courtship bouts approximated by IFD<8mm thresholding compared to manual scoring by a blinded observer. **b,c**, Comparison between the percent time courting and the average courtship bout length scored by IFD<8mm thresholding or scored by a blinded observer for the same courting pairs across species (b) or for *D. yakuba* male courtship of oenocyte-less females perfumed with the *D. yakuba* pheromone 7-T (c). Generally, IFD-thresholding overestimates the total amounts of courtship (due to incidental periods where flies are close but not courting) and underestimates the average courtship bout length (due to transient periods where the female moves away from the male but courtship continues), but captures the overall trends observed in larger samples manually scored by a blinded observer (d,e). **d**, Larger sample of manually scored courtship in the dark for conspecific courting pairs, *D. yakuba* males paired with a *D. simulans* females, or *D. simulans* males paired with a *D. yakuba* females. These heterospecific pairings suggest that differences between species in the dynamics of courtship in the dark are attributable to the males of the species rather than the females. **e**, Larger sample of manually scored courtship in the dark for *D. yakuba* male courtship of oenocyte-less females perfumed 7-T. Data points represent individual males and error bars are mean±SEM. Statistical tests performed were ANOVA with Tukey’s post-hoc (d) or unpaired Mann-Whitney (e). Letters above data sets denote statistically different groups (p<0.05). **** denote P<0.0001.

**Figure S2.**
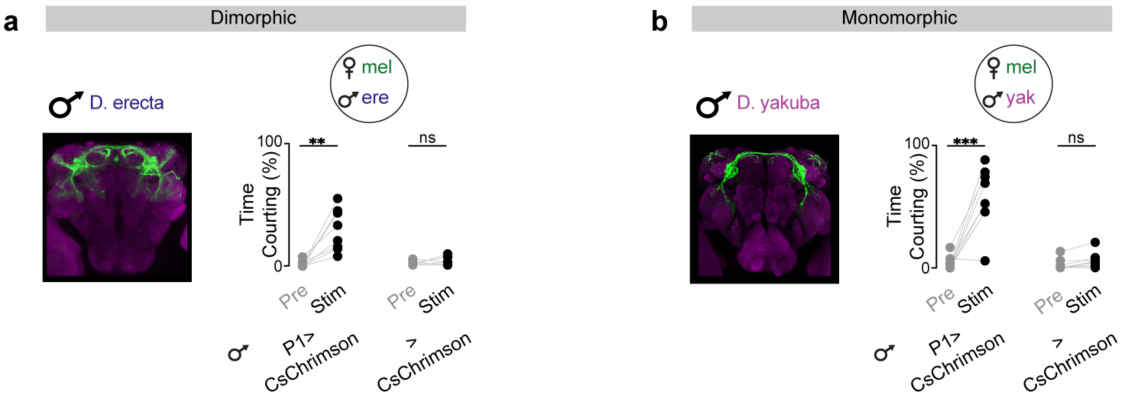
P1 neurons play a conserved role in promoting courtship across species. **a,b**, Percent time courting prior to (Pre) or during (Stim) periods of optogenetic stimulation of *D. erecta* (a) or *D. yakuba* (b) males expressing CsChrimson in P1 neurons (71G01>CsChrimson (a) or splitP1>CsChrimson (b)) or control animals (>CsChrimson) paired with a *D. melanogaster* female. Data points represent individual males. Significance was calculated by paired t-test, **P<0.01, ***P<0.001, ns, not significant.

**Figure S3.**
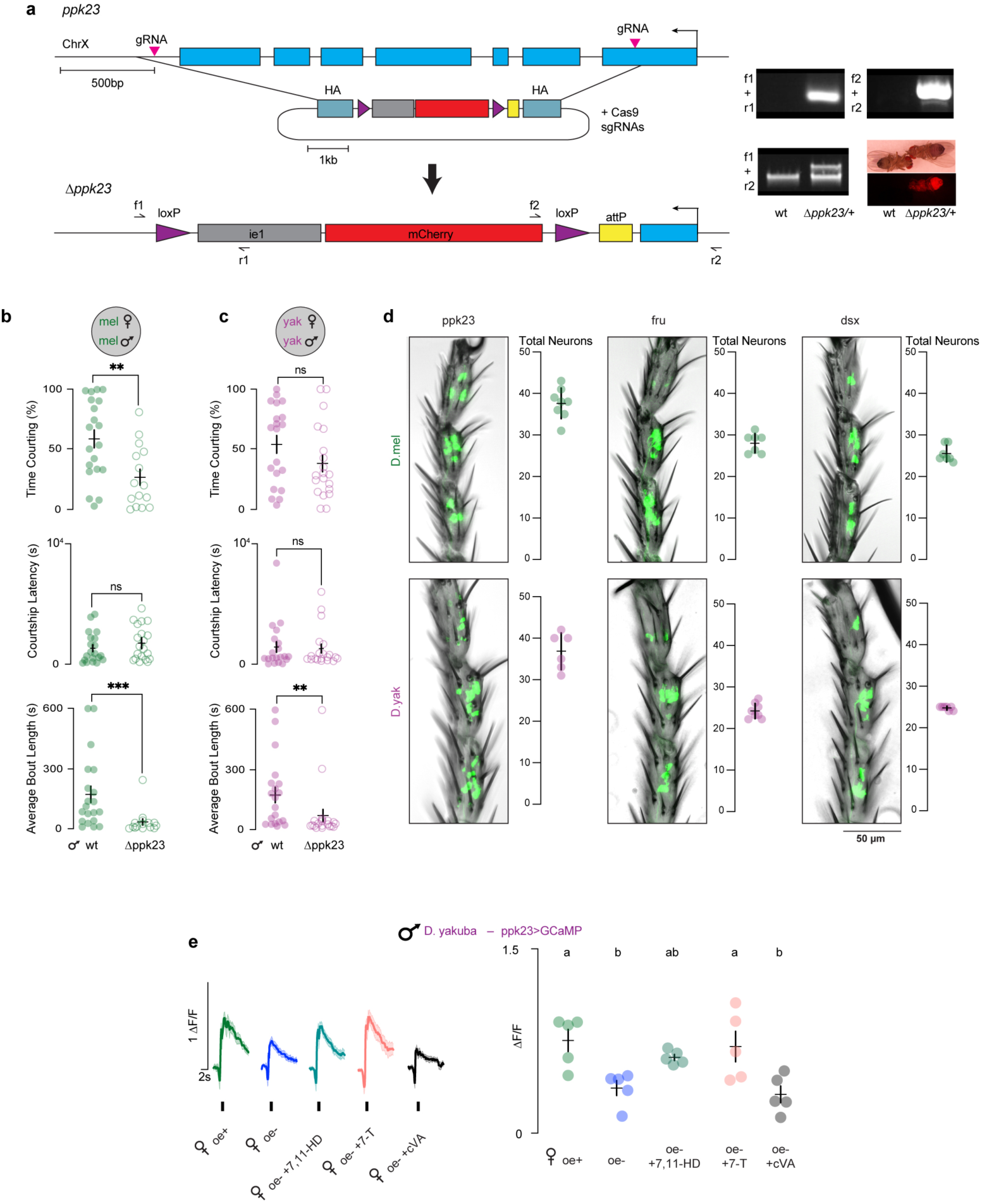
Conserved anatomy of pheromone-detecting sensory neurons. **a**, CRISPR/Cas9 targeting strategy for the genetic deletion of the *ppk23* locus and validation by PCR and ie1-mCherry marker expression. **b,c**, Percent time courting in the 10 minutes following courtship initiation, courtship latency, and courtship bout length of *ppk23* mutants relative to wildtype for *D. melanogaster* (b) and *D. yakuba* (c) males in the dark. **d**, Representative bright field images and quantification showing expression of CD8::GFP (green) in the soma of the Ppk23+, Fru+, and Dsx+ sensory neurons in the foreleg tarsal segments of *D. melanogaster* and *D. yakuba* males. Images are distal end up. **e,** (Left) Average functional responses (ΔF/F_0_) aligned to time of a tap (as in Fig. 3) of indicated *D. melanogaster* (oe+ ctrl), mock-perfumed oenocyte-less (oe-), 7,11-heptacosadiene-perfumed (oe- +7,11-HD), 7-tricosene-perfumed (oe- +7-T), or cis-Vaccenyl acetate-perfumed (oe- +cVA) female of the foreleg sensory afferents and (Right) average peak response (ΔF/F_0_) per male for a given female target. Data points represent individual males and error bars are mean±SEM. Statistical tests performed were ANOVA with Tukey’s post-hoc (e) or unpaired Mann-Whitney (b,c). Letters above data sets denote statistically different groups (p<0.05). Asterisks denote **P<0.01, ***P<0.001, ns, not significant.

**Figure S4.**
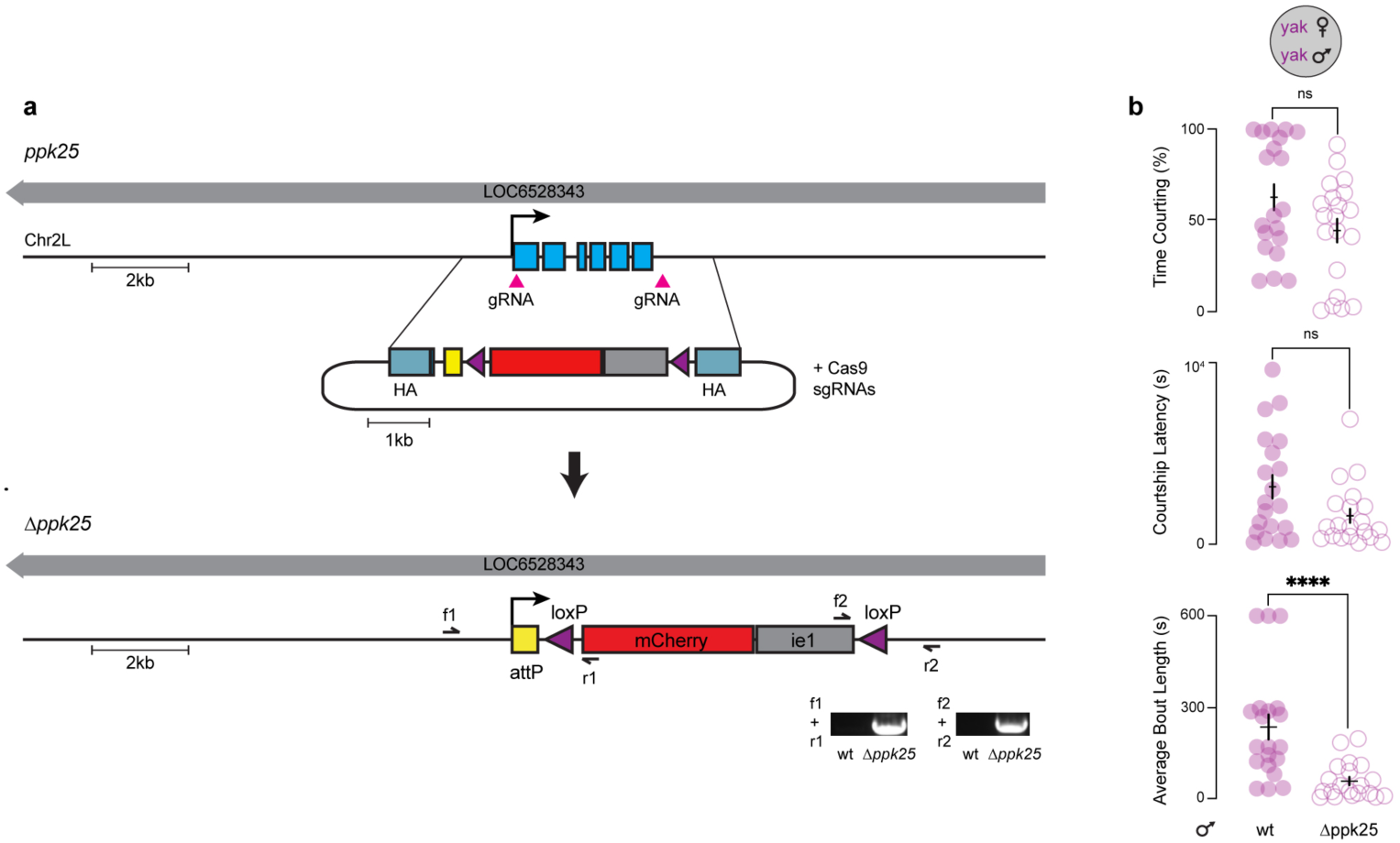
Ppk25+ Sensory neurons promote courtship in the dark in *D. yakuba*. **a,b**, CRISPR/Cas9 targeting strategy for the genetic deletion of the *ppk25* locus and validation by PCR (a) and courtship in the dark (as in Fig. S3) toward *D. yakuba* females (b). Shading in response traces represents mean±SEM. Data points represent individual males and error bars are mean±SEM. Significance was calculated by unpaired Mann-Whitney test. Asterisks denote ****P<0.0001, ns, not significant.

**Figure S5.**
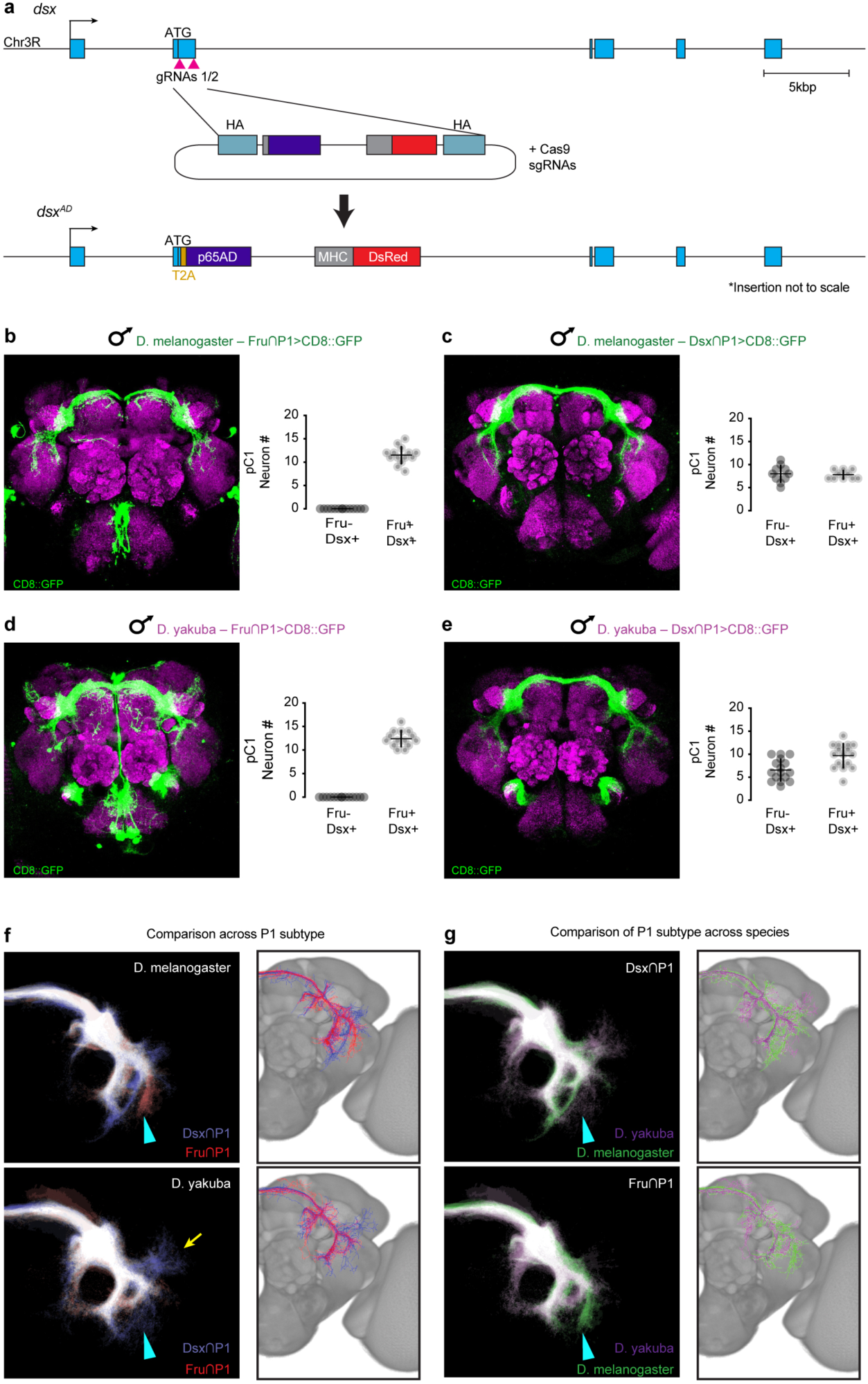
Differences in projection patterns of P1 subpopulations. **a**, CRISPR/Cas9 targeting strategy for the genetic insertion of a p65AD hemidriver into the *dsx* locus in *D. yakuba*. **b-e**, (Left) Unmasked images of Fru∩P1 and Dsx∩P1 neurons labeled by intersection of Fru (b,d) or Dsx (c,e) and the P1-driver 71G01 (Fru∩P1 and Dsx∩P1, respectively) in *D. melanogaster* and *D. yakuba*. Stained for GFP (green) and neuropil counterstain (magenta). (Right) Quantification of Dsx+ Fru∩P1 neurons and Fru^M^+ Dsx∩P1 neurons within the Dsx+ pC1 cluster, based on anti-Dsx and anti-Fru^M^ immunostaining, respectively. **f,g,** Comparison of the projections of Fru∩P1 and Dsx∩P1subpopulations across subtype (f) or across species (g). Heatmap (left) and skeletonized overlays (right) of the comparison groups. Regions of greatest difference across both subtype and species indicated (cyan triangle) and region of significant difference between Dsx∩P1 and Fru∩P1 in *D. yakuba* noted (yellow arrow). All images are dorsal side up.

**Figure S6.**
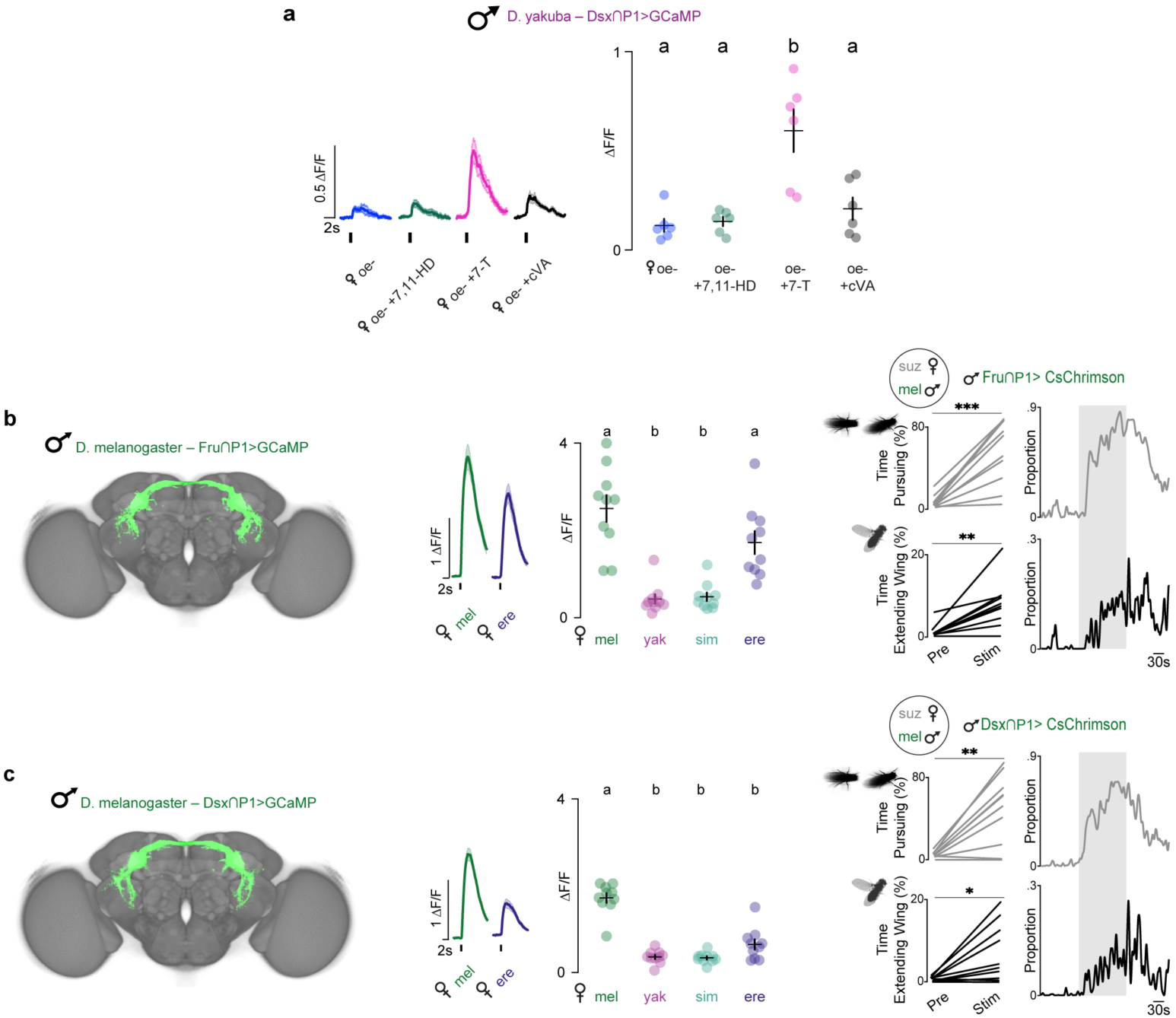
Subspecialization of P1 types in *D. melanogaster*. **a**, Averaged functional responses (ΔF/F_0_) of Dsx∩P1 in *D. yakuba* aligned to time of a tap of indicated *D. melanogaster* mock-perfumed oenocyte-less (oe-), 7,11-heptacosadiene-perfumed (oe- +7,11-HD), 7-tricosene-perfumed (oe- +7-T), or cis-Vaccenyl acetate-perfumed (oe- +cVA) female (left) and average peak response (ΔF/F_0_) per male for a given female target (right). **b,c, (**Left) Maximum intensity projections of neurons labeled by intersection of Fru (a) or Dsx (b) and the P1-driver 71G01 (Fru∩P1 and Dsx∩P1, respectively) in *D. melanogaster*, registered to a common template brain (see methods). (Middle) Averaged functional responses (ΔF/F_0_) aligned to time of a tap of indicated conspecific or heterospecific female and average peak response (ΔF/F_0_) per male for a given female target. (Right) Total percent time pursuing (top) or performing unilateral wing extensions (bottom) toward a *D. suzukii* (suz) female target prior to (Pre) or during (Stim) optogenetic stimulation of Fru∩P1>CsChrimson or Dsx∩P1>CsChrimson males and the proportion of flies engaging in these behaviors over the course of the experiment. Shading in response traces represents mean±SEM. Data points represent individual males and error bars are mean±SEM. Statistical tests performed were ANOVA with Tukey’s post-hoc (a-c) or paired t-test (b,c). Letters above data sets denote statistically different groups (p<0.05). Asterisks denote *P<0.05, **P<0.01, ***P<0.001.

**Figure S7.**
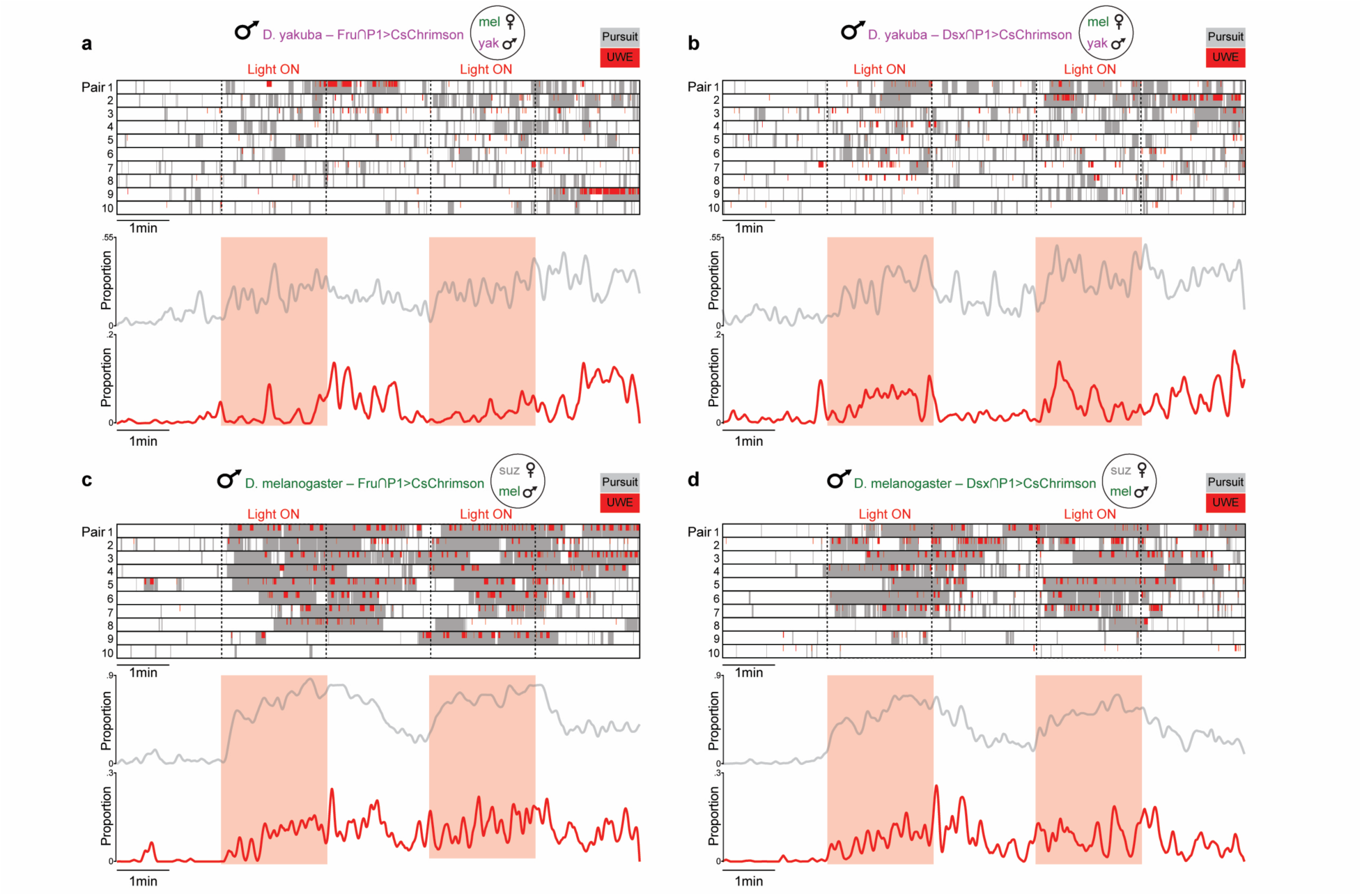
Activation of P1 subtypes. **a-d,** (Top) Raster plots of female pursuit (gray) or unilateral wing extension behaviors (UWE; red) exhibited upon optogenetic stimulation of Fru∩P1>CsChrimson (a,c) or Dsx∩P1>CsChrimson (b,d) males and the proportion of flies engaging in these behaviors over the course of the experiment in *D. yakuba* (a,b) or *D. melanogaster* (c,d). Dotted block lines and areas of red shading indicate stimulation.

**Figure S8.**
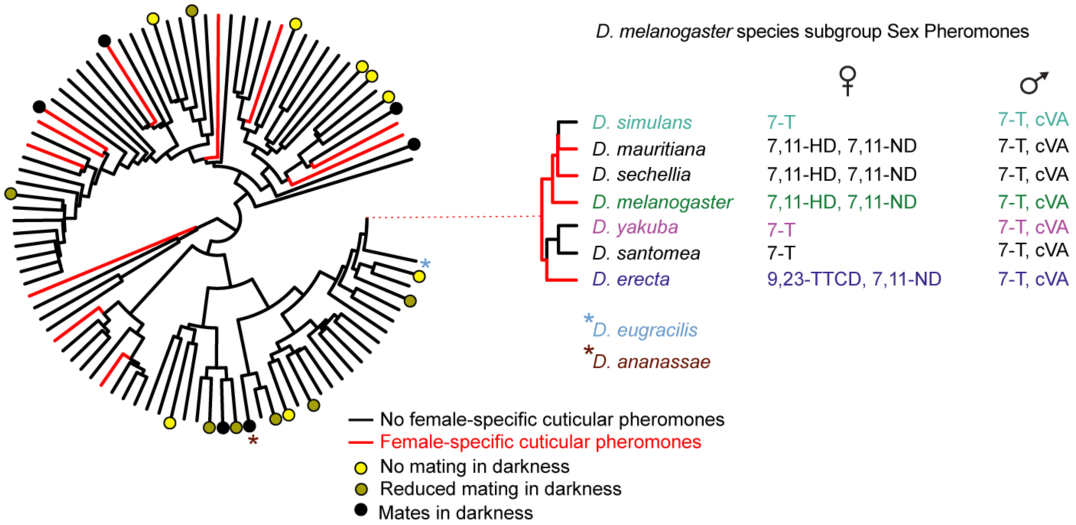
Mating in the dark has arisen repeatedly in monomorphic species. Phylogeny of Drosophila species (as in Fig. 1) with previous reports of various species’ capacity to mate in the dark indicated (*42*, *43*, *60–64*).

**Figure S9:**
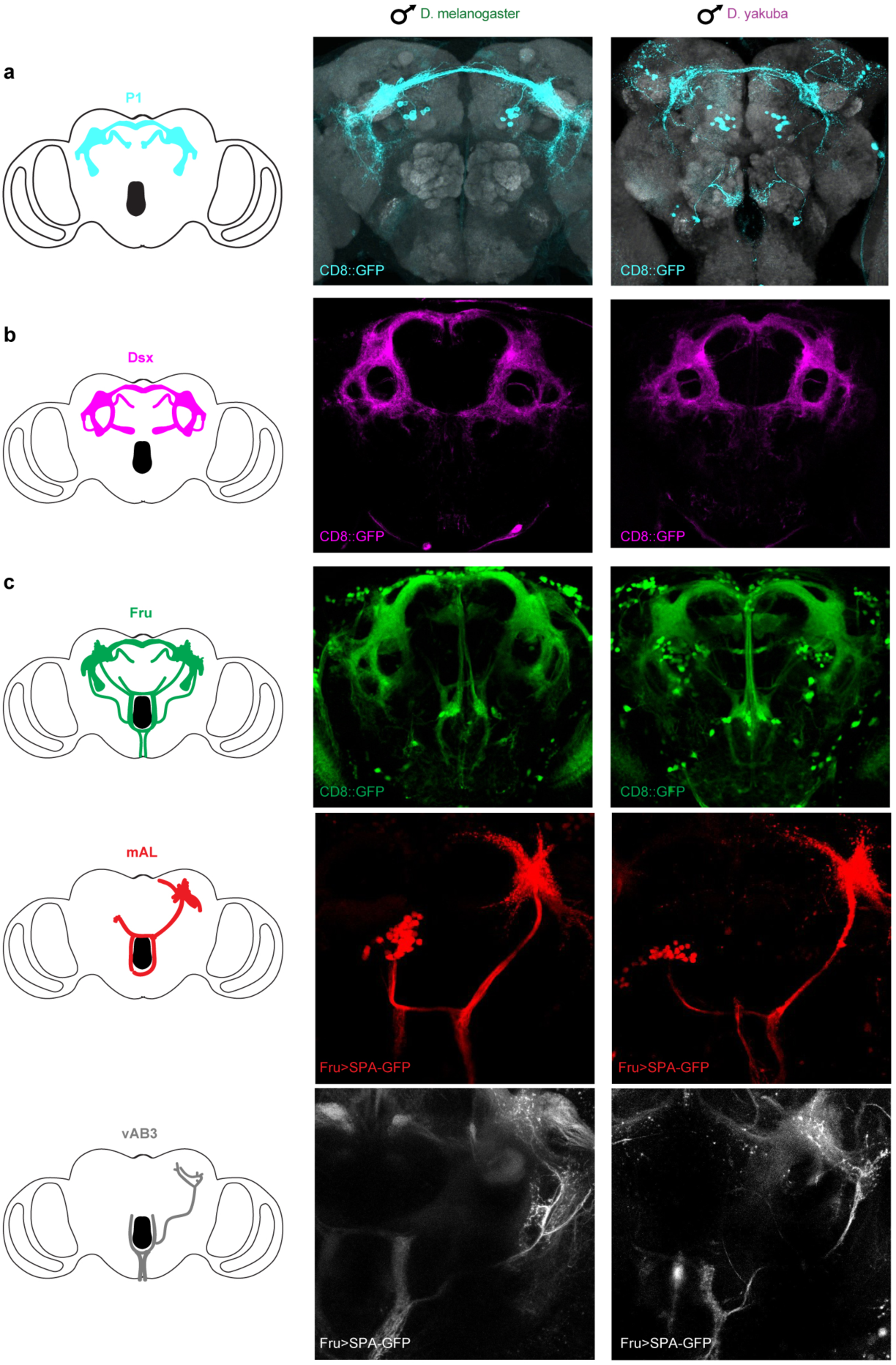
Overall anatomical conservation of the Fru+, Dsx+, and ascending circuitry. **a**, Unmasked splitP1>CD8::GFP expression in the brains of *D. melanogaster* (left) and *D. yakuba* (right) males. Stained for GFP (green) and neuropil counterstain (gray). **b**, Dsx>CD8::GFP (green) expression in the brains of *D. melanogaster* (left) and *D. yakuba* (right) males. **c**, (Top) Fru>CD8::GFP expression, as in (b). (Middle and bottom) Photoactivation labeling of mAL and vAB3, respectively, using Fru>SPA-GFP. All images are dorsal side up.

## Methods

### Fly stocks and husbandry

Flies were maintained at 25°C and 50–65% relative humidity under a 12 hr light:12 hr dark cycle. *D. melanogaster* stocks Canton-S (64349), UAS-GCaMP6s (42746 and 42749), UAS-mCD8-GFP (BL# 5137), 71G01-Gal4 (39599), fru^LexA^ (66698), and UAS-CsChrimson-mVenus (55134 and 55136), were obtained from the Bloomington Stock Center. LexAop-SPA-T2A-SPA was generated in a previous study(*25*). *D. yakuba* Ivory Coast (14021-0261.00), *D. erecta* (14021-0224.01), and *D. ananassae* (14024-0371.34) were obtained from the Cornell (formerly UCSD) Stock Center; D. eugracilis (SHL12) was obtained from the Kyoto Stock Center; D. suzukii (WT3) (L. Zhao, Rockefeller). *D. melanogaster* SplitP1-Gal4 (D. Anderson, Caltech); *D. melanogaster* fru^Gal4^ and fru^AD^/fru^DBD^ (B. Dickson, Janelia Research Campus); *D. melanogaster* ppk23-Gal4 (K. Scott, UC Berkeley); *D. yakuba* fru^Gal4^, fru^DBD^, fru^AD^ 71G01-Gal4, UAS-GCaMP6s, UAS-CsChrimson, UAS-Kir2.1 and attp1730 (D. Stern, Janelia Research Campus). *D. erecta* 71G01-Gal4 (insertion #5), *D. erecta* UAS-GCaMP6s (insertion #5), *D. erecta* UAS-CD8::GFP (insertion #2), *D. erecta* UAS-CsChrimson::tdTomato (insertion #1), *D. yakuba* dsx^Gal4^ and dsx^AD^ (Y. Ding, University of Pennsylvania), *D. yakuba* UAS-mCD8-GFP (insertions #2 & 3), UAS-GFP (insertion #8), UAS-SPA-GFP (insertion #2), ppk23-Gal4 (insertion #2), *ppk23* mutant, and *ppk25* mutant were generated in this study. See Supplemental Table 1 for detailed genotypes by figure.

### Construct design and generation

The ie1.mCherry and ie1.eGFP minigene reporters used as transformation markers were constructed from the following fragments: the ie1 promoter PCR amplified from the pBac{orco.QF, QUAS.GCaMP, ie1. DsRed} (gift from D. Kronauer), *Drosophila* codon-optimized mCherry and eGFP gene blocks (IDT), and the p10-3’UTR PCR amplified from the plasmid pJFRC81 (Addgene #36432). These were then cloned by Gibson assembly (NEB) in to the backbone of pBac{3XP3-EGFP, Pactin-Ptrsps} (Addgene #86861) after digest with BstZ17I and Bsu36I to generate pRC7a (ie1.mCherry) and pRC10a (ie.eGFP).

To create a more flexible multicloning site in pRC10a a gBlock (IDT) containing the following restriction enzyme sites HpaI, HincII, HindIII, KpnI, NotI, XhoI, XbaI and FseI was synthesized and cloned by Gibson assembly into pRC10a. The resulting vector was pIM145. mCherry was then cloned NheI and HpaI from pRC7a into the pIM145 backbone in place of GFP to generate pIM148.

To generate a P1 driver line, the 3844 base pair fragment from Vsx2 gene that is present pGMR71G01, was amplified from *D. melanogaster* genomic DNA. This was then cloned by Gibson assembly into the backbone of pIM145 in front of a Gal4 gene block (IDT), generating pRC12b.

To generate a BAC vector containing UAS-GFP the following cassette 10X UAS-IVS-Syn21-GFP p10 UTR was excised from pJFRC81 (Addgene 36432) with HindIII and FseI and ligated into pIM145. The resulting vector was pIM153.

To generate a BAC vector containing UAS-SPA-GFP a *Drosophila* codon optimized SPA-GFP was synthesized as a gBlock (IDT) and cloned by Gibson assembly (NEB) into pIM147 in place of GCaMP6 to generated pIM141.

To generate ppk23-Gal4 a DNeasy Blood & Tissue kit (Qiagen) was used to purify wild type *D. yakuba* genomic DNA. 50 mg of flies were collected and snap frozen in liquid nitrogen. Flies were ground to a fine powder using a mortar and pestle. Powder was transferred to a microcentrifuge tube and processed according to the DNeasy Blood & Tissue kit instructions. The resulting genomic DNA was used as a template for PCR amplification of the ppk23 promoter. The 2.7 KB promoter was cloned into pRC12B to generate pIM169.

To generate a BAC vector containing a blue fluorescent marker for visualization a *Drosophila* codon optimized mTagBFP2 was synthesized. Due to the sequence complexity of mTagBFP2 it was designed as two overlapping gBlocks (IDT) and included a portion of the ie1 promoter for seamless assembly into the destination vector. mTagBFP2 was cloned by Gibson assembly (NEB) into pIM148 in place of mCherry to generated pIM149.

### CRISPR-Cas9 mediated deletion of *ppk23* in *D. yakuba*

Two gRNAs were designed to direct Cas9-mediated cleavage to the first exon and 3’ UTR of the *ppk23* gene locus in *D. yakuba*. gRNA off-target potential was determined using CRISPR optimal target finder (http://tools.flycrispr.molbio.wisc.edu/targetFinder/index.php) gRNA sequences, gRNA1 CATCGGTGCGGTCACCGCAC and gRNA2 GTGTTGCATACTTAGCGGCG, were PCR amplified with Q5 High-Fidelity master mix (NEB) and cloned into pCFD4 (Addgene 49411) by Gibson assembly (NEB). The resulting vector was pIM179. pIM148 was digested HpaI and SmaI to liberate ie1p mCherry p10 UTR. The mCherry cassette was then ligated to pDsRed-attP (Addgene 51019) which was digested with AgeI and BsiWI to remove the 3xP3-DsRed cassette then Klenow endfilled. This generated pIM174. 1 Kb homology arms beginning at the predicted Cas9 cut sites in *ppk23* were PCR amplified with Q5 High-Fidelity master mix (NEB) and cloned into pIM174. The resulting vector was pIM175. A cocktail of pIM179, pIM175 and Cas9 protein was injected into wild type *D. yakuba* embryos by Rainbow Transgenic Flies using standard injection procedures. Viable G0 flies were mated to wild type male or virgin female flies. F1 progeny were screened visually by mCherry expression. mCherry positive F1s were individually crossed to wild type male or virgin female flies then sacrificed for genomic DNA. Deletion of *ppk23* was confirmed in mCherry positive F1s by genotyping using primers internal to and flanking the targeted genomic region. PCR products from genotyping were sequenced to verify exact genome modification. mCherry positive F2s from sequence verified F1s were self-mated and mCherry positive F3 virgin females were genotyped by nonlethal methods to identify females homozygous for *ppk23* deletion. As the *ppk23* locus is on the X chromosome, virgin females homozygous for the *ppk23* null mutation were then mated to males hemizygous for the mutation to produce a stable line.

### CRISPR-Cas9 mediated deletion of *ppk25* in *D. yakuba*

Two gRNAs were designed to direct Cas9-mediated cleavage to the first exon and 3’ UTR of the ppk25 gene locus in D. yakuba. gRNA off-target potential was determined using CRISPR optimal target finder (http://tools.flycrispr.molbio.wisc.edu/targetFinder/index.php) gRNA sequences, gRNA1 GUCGGUCGAUGCAACCGGAC and gRNA2 UAAACUUAACAACAUCGGAG were synthesized from Synthego. 1 Kb homology arms beginning at the predicted Cas9 cut sites in ppk25 were ordered as gBlocks from IDT. The ppk25 start code in the 5’ homology arm was mutated from ATG to TTG. The homology arms were cloned sequentially into pIM174 by Gibson assembly. The resulting vector was pIM188. A cocktail of pIM188, sgRNA1, sgRNA2 and Cas9 protein was injected into wild type D. yakuba embryos by Rainbow Transgenic Flies using standard injection procedures. Viable G0 flies were mated to wild type male or virgin female flies. F1 progeny were screened visually by mCherry expression. mCherry positive F1s were individually crossed to wild type male or virgin female flies then sacrificed for genomic DNA. Deletion of ppk25 was confirmed in mCherry positive F1s by genotyping using primers internal to and flanking the targeted genomic region. PCR products from genotyping were sequenced to verify exact genome modification. mCherry positive F2s from sequence verified F1s were self-mated and mCherry positive F3 virgin females and males were genotyped by nonlethal methods to identify flies homozygous for ppk25 deletion.

### CRISPR-Cas9 mediated insertion into of *dsx* in *D. yakuba*

For the T2A-Gal4 insertion into *dsx*, a pair of gRNAs were designed using DRSC CRISPR finder (https://www.flyrnai.org/crispr/). gRNA1 (GGAGTCGATCATGTCCGAGT; first G was added to enhance transcriptional efficiency) and gRNA2 (GGCGCCGTTCTTGCGATTGA), were PCR amplified using Phusion High-Fidelity PCR Master Mix (NEB) and cloned into pCFD4 (Addgene 49411) by Gibson assembly (NEB). The *dsx* T2A-GAL4 knock-in allele was generated by replacing the majority of the first coding exon (second exon) of dsx in-frame (39bp from ATG) with T2A-GAL4. The donor plasmids were constructed by concatenating a 1.2 kb left homology arm, T2A, GAL4 (amplified from pBPGUw, Addgene #17575), a Mhc-DsRed marker, a 1.0 kb right homology arm, and a 1.8 kb backbone using Gibson Assembly (NEB). T2A was incorporated into the plasmid by using a gBlock (IDT) that covers the left homology arm and T2A. A cocktail of pCFD4 gRNA plasmid, donor plasmid, and cas9 mRNA was injected into wild type D. yakuba embryos by Rainbow Transgenic Flies using standard injection procedures. F1 progeny was screened for MhC-DsRed, which drives red fluorescence expression in muscles. Transformants were verified by PCR genotyping and maintained through screening for flies carrying Mhc-DsRed. The T2A-p65-AD insertion into *dsx* was generated using the same method but a different gRNA2 (GGCGGACCGCCAGCGGGTGA). The p65-AD sequence was amplified from pBPp65ADZpUw (addgene 26234) before Gibson assembly as above.

### Immunohistochemistry

Adult brains were dissected in Schneider’s media (Sigma) then immediately transferred to cold 1% PFA (Electron Microscopy Sciences) and fixed overnight at 4°C. Following overnight incubation samples were washed in PAT3 Buffer (0.5% BSA/0.5% Triton/1X PBS pH 7.4) 3 times. Brains were blocked in 3% Normal Goat Serum for 90 minutes at RT. Primary antibodies 1:1000 chicken anti-GFP (Abcam ab13970), 1:50 mouse anti-brp (Developmental Studies Hybridoma Bank nc82), 1:2000 rabbit anti-Fru^M^ (generated for this study by YenZyme against a synthesized peptide: HYAALDLQTPHKRNIETDV^70^) and 1:500 guinea pig anti-Fru^M,^ (gift from D. Yamamoto, Tohoku University) were incubated 3 hours at RT then 2-3 days at 4°C. Brains were washed extensively in PAT3 Buffer. Secondary Alexa Fluor antibodies (Life Technologies) were incubated 3 hours at RT then 2-3 days at 4°C. Brains were washed 3 times in PAT3 Buffer then once in 1X PBS. Samples were mounted in Vectashield (Vector Laboratories). Images were captured on a Zeiss LSM 880 using a Plan-Apochromat 20X (0.8 NA) objective.

Leg images were taken using the native fluorescence of animals expressing UAS.CD8::GFP (insertion #2). Animals were aged ∼3-5 days old and legs were mounted in Vectashield and femurs stabilized using UV glue. Images were taken at 25x with 1.6x digital zoom.

For analysis of P1 projections, images were registered to the JRC2018M template brain using Computational Morphometry Toolkit (https://www.nitrc.org/projects/cmtk/) and P1 neurons were segmented in VVD Viewer (https://github.com/JaneliaSciComp/VVDViewer).

### Courtship assays in the light

All single choice assays were conducted at 25°C, 50-65% relative humidity between 0 to 3 hours after lights on. Male flies were collected shortly after eclosion and group housed for 4-7 days prior to assay. Target virgin females were 4-7 days post-eclosion. Assays were performed in 38 mm diameter, 3 mm height circular chambers in a 4 × 4 array back lit using a white light pad (Logan Electric). Fly behavior was recorded from above the chambers using a PowerShot SX620 camera (Canon) or Point Grey FLIR Grasshopper USB3 camera (GS3-U3-23S6M-C: 2.3 MP, 162 FPS, Sony IMX174, Monochrome). A virgin female was transferred to the chamber by mouth aspiration followed by a test male. Once the male was loaded into the chamber the assay commenced and the activity of the flies was recorded for 10 minutes. Orienting, wing extension, chasing, mounting and copulation were used to score courtship behavior during video analysis by a blinded observer. Note that courtship assays in the light were not conducted on food to avoid flies copulating too quickly under these conditions. In the absence of food, females generally move much quicker. This results in shorter male courtship bouts, likely explain the difference in bout lengths observed between males in the dark on food compared to males in the light off food (i.e. bouts were generally longer in the dark when on food).

### Courtship assays in the dark

Courtship assays performed in the dark were conducted at 25°C, 50-65% relative humidity between 0 to 3 hours after lights on. Male flies were collected shortly after eclosion and group housed for 4-7 days prior to assay. Target virgin females were 4-7 days old. Assays were performed on food in 35mm x 10mm Petri Dishes (Falcon). Females were transferred to the chamber by mouth aspiration followed by a test male. Once the male was loaded into the chamber the assay commenced and the activity of the flies was recorded for 3 hours. Assays were back-lit by infrared LED strips (940 nm, LED Lights World). Fly behavior was recorded from above the chambers using a Point Grey FLIR Grasshopper USB3 camera (GS3-U3-23S6M-C: 2.3 MP, 162 FPS, Sony IMX174, Monochrome) using the Flycapture2 Software Development Kit (FLIR). Orienting, wing extension, chasing, mounting and copulation were used to score courtship behavior during video analysis by an observer blind to the genotype or species and inter-fly distance was quantified using FlyTracker (Caltech). Unfortunately, variations in the food and/or infrared lighting meant that for approximately half of the videos collected we could not obtain high-quality tracking data. Thus, we opted to track a random subset, i.e. the first six consecutive courting pairs from which quality tracking was possible. This provides a representative visual assessment of courtship dynamics, while the complete data set of all courting pairs was scored manually by blinded observer. Fig. S1 validates that while IFD-tracking and manually scored datasets differed in the absolute quantity of courtship scored, the overall trends in the data were consistent.

### Generation of oenocyteless animals

Oenocyte ablation was performed genetically by crossing male +;PromE(800)-Gal4, tub-Gal80^TS^;+ flies to female +;UAS-StingerII, UAS-hid/CyO;+ at 18 °C. Newly eclosed virgin females were collected and kept at 25 °C for 1 day. Females were then shifted to 30 °C for 2 days and then allowed to recover at 25 °C for 2 days before use in experiments. Females were screened for GFP expression to confirm oenocyte ablation.

### Perfuming

Mock, 7-T, and, 7,11-HD perfuming of oenocytel-less or *D. yakuba* females was performed by filling adding heptane, 2µg 7-T, or 2µg 7,11-HD (Cayman Chemicals), respectively, to 1mL of heptane in a 32mm scintillation vial (ThermoScientific). A vacuum was then applied to evaporate the heptane from the vial. 5-8 flies were added to each vial and vortexed at low speed for 30 seconds, 3 times. cVA perfuming was perfumed by dissolving cVA to a concentration of 5mg/mL in ethanol. 0.5µL was then applied directly to each fly’s abdomen by pipette. All flies were returned to food to recover for at least 1 hour prior to experimentation.

### Photoactivation

For photoactivation experiments used fru^Gal4^ expressed SPA-GFP. Photoactivation was performed on adult flies aged 24–48 h after eclosion. Brains were imaged at 925nm to identify suitable sites for photolabeling while not stimulating photoconversion. Using PrairieView, an ROI was then drawn around projections unique to the cell type of interest and photoconversion was stimulated in a single z-plane by brief exposure of the ROI to 710nm laser light. Power was 5–35 mW at the back aperture of the objective, depending on the depth of the neurite being converted. This process was repeated 50-100 times, interposed with rests to allow for diffusion of the photoconverted molecules, until the cell type of interest was uniformly above background levels of fluorescence.

### Functional imaging

All imaging experiments were performed on an Ultima two-photon laser scanning microscope (Bruker) equipped with galvanometers driving a Chameleon Ultra II Ti:Sapphire laser. Emitted fluorescence was detected with a GaAsP photodiode (Hamamatsu) detectors. All images were collected using PrairieView Software at 512 pixel × 512 pixel resolution. VNC and LPC preparations were performed as previously reported (*17*, *25*)

For the *D. yakuba* P1 imaging, “splitP1” (71G01-AD 15A01-DBD intersection) had to be used rather than 71G01-Gal4 because this line was weak and bleached before an experiment could be completed. These splitP1 animals showed inter-animal variability in responses. Only ∼1 in 5 (5/27) animals exhibited P1 responses to conspecifics, though in responding animals the responses were uniform and predictably evoked each time a male tapped a conspecific female (as shown in Fig. 2). We discovered this likely resulted from stochastic labeling of Fru-Dsx+ P1 subset. Specifically, splitP1 predominantly labels Fru+ P1 neurons, and labels 2-3 Fru-Dsx+ neurons in only some animals, as previously found in D. melanogaster (*57*). Indeed, when we subsequently imaged Dsx∩P1 (71G01-DBD dsx-AD intersection) no inter-animal variability was observed and all animals responded to conspecifics with each tap. The same was true when imaging for all Dsx+ projections in the LPC. Therefore, for simplicity and understanding we have presented just responding *D. yakuba* splitP1 males in Fig. 2b.

### Optogenetic set-up

Optogenetic assays were performed in a 38 mm diameter, 3 mm height circular chamber with sloping walls. The chamber was placed in the middle of a 3-mm thick acrylic sheet suspended on aluminum posts above a 3 × 4 array of 627 nm LEDs (Luxeon Star LEDs). LEDs were attached to metal heat sinks (Mouser Electronics) which were secured at 5 cm intervals to a 30 × 30 cm aluminum wire cloth sheet (McMaster-Carr). LEDs were driven by Recom Power RCD-24-0.70/W/X2 drivers, which were powered by a variable DC power supply. Infrared LED strips (940 nm, LED Lights World) attached to the wire cloth between the heat sinks provided back-illumination of the platform. LED strips were covered with 071 Tokyo blue filter (Lee Filters) to remove potential activating light emitted from the illumination source. LED drivers were controlled by the output pins of an Arduino running custom software. Fly behavior was monitored from above the chamber using a Point Grey FLIR Grasshopper USB3 camera (GS3-U3-23S6M-C: 2.3 MP, 162 FPS, Sony IMX174, Monochrome) outfitted with 071 Tokyo blue filter (Lee Filters) to avoid detection of light from the high-power LEDs. Flies were recorded at 30 frames/sec. Custom software was used for data acquisition and instrument control during assays. Light intensity was measured with a photodiode power sensor (Coherent 1212310) placed at the location of the behavioral chamber. The peak wavelength of the LED (627 nm) was measured across a range of voltage inputs. Measurements were repeated three times and averaged. The baseline intensity before LED illumination was subtracted.

### Optogenetic assays

Flies were reared on standard SY food in the dark at 25°C and 50-65% relative humidity. *P1>UAS-CsChrimson* and UAS-CsChrimson control male flies were collected shortly after eclosion, group housed for 3 days then shifted to food containing 0.4 mM all trans-retinal (Sigma) for 48 hours before being assayed. ppk23-Gal4>UAS-CsChrimson and UAS-CsChrimson control male flies were collected shortly after eclosion, group housed for 3 days then shifted to food containing 0.4 mM all trans-retinal for 24 hours before being assayed. Fru∩P1>UAS-CsChrimson, Dsx∩P1>UAS-CsChrimson, and UAS-CsChrimson control males were collected shortly after eclosion, grouped housed for 2 days and then shifted to food containing 0.4 mM all trans-retinal for 48 hours before being assayed. Target virgin females were 4-6 days old. Flies were added to a 38 mm diameter, 3 mm height circular courtship chamber by mouth aspiration. Once a male was transferred to a courtship chamber containing a virgin female the assay commenced and the activity of the flies was recorded for 10 minutes. Neurons expressing CsChrimson were activated by 627 nm wavelength stimulation. To activate splitP1 (w; 71G01-AD/+; 15A01-DBD/UAS-CsChrimson) neurons in *D. yakuba* the following stimulation protocol was used: 2 minutes dim white light followed by 2 minutes 627 nm LED (5 Hz, 100 ms pulse-width, 10 µW/mm^2^) alternating for 10 minutes total. For activation of P1 (UAS-CsChrimson::tdTomato /+; 71G01-Gal4/+) neurons in *D. erecta* the stimulation protocol was: 2 minutes dim white light followed by 2 minutes 627 nm LED (5 Hz, 100 ms pulse-width, 3.4 µW/mm^2^) alternating for 10 minutes total. To activate Ppk23+ neurons in *D. yakuba* the following stimulation protocol was used: 2 minutes dim white light followed by 2 minutes 627 nm LED (5 Hz, 100 ms pulse-width, 8 µW/mm^2^) alternating for 10 minutes total. All assays were conducted at 25°C, 50-65% relative humidity between 0 to 3 hours after lights on. To activate Fru∩P1 and Dsx∩P1 neurons in *D. yakuba* the following stimulation protocol was used: 2 minutes dim white light followed by 2 minutes 627 nm LED (5 Hz, 100 ms pulse-width, 10 µW/mm^2^) alternating for 10 minutes total. The same stimulation protocol was used in *D. melanogaster* males for each P1 populations, except the light intensity used was 3.4 µW/mm^2^. A blinded observer scored orienting, wing extension, chasing, mounting and copulation to quantify courtship behavior for the purposes of calculating overall courtship indices. To quantify dynamic behaviors, videos were tracked using FlyTracker (Caltech). Behavioral classifiers for courtship pursuit behavior (consisting or orienting toward the female and moving toward her) and unilateral wing extensions were then trained using JAABA (Janelia) in each species.

### Statistical analysis

Statistical analyses was performed in GraphPad Prism 9. Before analysis, normality was tested for using the Shapiro–Wilk method for determining whether parametric or non-parametric statistical tests would be used. In cases where multiple comparisons were made, appropriate post-hoc tests were conducted as indicated in figure legends. All statistical tests used were two-tailed. Experimenters were blind to experimental conditions during analysis. See Supplemental Table 2 for detailed statistics by figure.

---

**Table S1.** Male experimental genotypes by figure.

| Figure | Species | Male Genotype |
| --- | --- | --- |
| 1b-d; 3a,d,e,f; S1a-e; S3c; S4c | *D. yakuba* | Wildtype (Ivory Coast) |
| 1b,c; 3c; S1b; S3b; | *D. melanogaster* | Wildtype (Canton S) |
| 1b,c; 2b; 3e; 4a-e; S1a,b,d; S6b,c | *D. simulans* | Wildtype |
| 1b,c; 2b; 3e; 4a-e; S6b,c | *D. erecta* | Wildtype (14021-0224.01) |
| 1b,c | *D. eugracilis* | Wildtype (SHL12) |
| 1b,c | *D. ananassae* | Wildtype (14024-0371.34) |
| 2a,b | *D. melanogaster* | w; 71G01-Gal4/UAS-CD8::GFP |
| 2a,b | *D. melanogaster* | w; UAS-GCaMP6s/+; 71G01-Gal4/+ |
| 2a,b | *D. erecta* | UAS-CD8::GFP/+; 71G01-Gal4/+ |
| 2a,b | *D. erecta* | UAS-GCaMP6s/+; 71G01-Gal4/+ |
| 2a,b | *D. simulans* | w; UAS-GCaMP6s/+; 71G01-Gal4/+ |
| 2a,b | *D. yakuba* | w; 71G01-AD/+; 15A01-DBD/UAS-CD8::GFP |
| 2a,b | *D. yakuba* | w; 71G01-AD/UAS-GCaMP6s; 15A01-DBD/+ |
| 3a; S3d | *D. yakuba* | w; ppk23-Gal4/UAS-CD8::GFP |
| 3a; S3d | *D. melanogaster* | w; ppk23-Gal4/+; UAS-CD8::GFP/+ |
| 3c; S3b | *D. melanogaster* | Δppk23 |
| 3b,d; S3c | *D. yakuba* | Δppk23 |
| 3e; S4a | *D. yakuba* | w; UAS-GCaMP6s; ppk23-Gal4/+ |
| 3e | *D. melanogaster* | w; ppk23-Gal4; UAS-GCaMP6s |
| 3e | *D. yakuba* | Δppk23; UAS-GCaMP6s/+; ppk23-Gal4/+ |
| 3e | *D. yakuba* | w; UAS-GCaMP6s, Δppk25/Δppk25; fru-Gal4/+ |
| 3f; S4c | *D. yakuba* | Δppk25 |
| 3g | *D. yakuba* | w; ppk23-Gal4/UAS-CsChrimson |
| 3g | *D. yakuba* | w; UAS-CsChrimson/+ |
| 4a; S5d,f,g | *D. yakuba* | w; 71G01-DBD/+; fru-p65.AD/UAS-CD8::GFP |
| 4b; S5e-g | *D. yakuba* | w; 71G01-DBD/+; dsx-p65.AD/UAS-CD8::GFP |
| 4a | *D. yakuba* | w; 71G01-DBD/UAS-GCaMP6s; fru-p65.AD/+ |
| 4b; S6a | *D. yakuba* | w; 71G01-DBD/UAS-GCaMP6s; dsx-p65.AD/+ |
| 4a; S7a | *D. yakuba* | w; 71G01-DBD/+; UAS-CsChrimson, fru-p65.AD/+ |
| 4b; S7b | *D. yakuba* | w; 71G01-DBD/+; UAS-CsChrimson, dsx-p65.AD/+ |
| 4c | *D. yakuba* | w; UAS-GCaMP6s/UAS-GCaMP6s; fru-Gal4/+ |
| 4d | *D. yakuba* | w; UAS-GCaMP6s/UAS-GCaMP6s; dsx-Gal4/+ |
| 4e | *D. yakuba* | Δppk23; UAS-GCaMP6s/+; dsx-Gal4/+ |
| 4f | *D. yakuba* | w; 71G01-DBD/+; UAS-Kir2.1/+ |
| 4g | *D. yakuba* | w; 71G01-DBD/+; fru-p65.AD/UAS-Kir2.1 |
| 4h | *D. yakuba* | w; 71G01-DBD/+; dsx-p65.AD/UAS_Kir2.1 |
| S2a | *D. erecta* | UAS-CsChrimson::tdTomato /+; 71G01-Gal4/+ |
| S2b | *D. yakuba* | w; 71G01-AD/+; 15A01-DBD/UAS-CsChrimson |
| S3d | *D. melanogaster* | w; fru-Gal4/UAS-CD8::GFP |
| S3d | *D. yakuba* | w; fru-Gal4/UAS-CD8::GFP |
| S3d | *D. melanogaster* | w; dsx-Gal4/UAS-CD8::GFP |
| S3d | *D. yakuba* | w; dsx-Gal4/UAS-CD8::GFP |
| S5b,f,g; S6b | *D. melanogaster* | w; 71G01-p65.AD/+; UAS-CD8::GFP, fru-DBD/+ |
| S5c,f,g; S6c | *D. melanogaster* | w; 71G01-p65.AD/UAS-CD8::GFP; dsx-DBD/+ |
| S6b | *D. melanogaster* | w; 71G01-p65.AD/+; UAS-GCaMP6s, fru-DBD/+ + |
| S6c | *D. melanogaster* | w; 71G01-p65.AD/UAS-GCaMP6s; dsx-DBD/+ |
| S6b; S7c | *D. melanogaster* | w; 71G01-p65.AD/+; UAS-CsChrimson, fru-DBD/+ + |
| S6c; S7d | *D. melanogaster* | w; 71G01-p65.AD/UAS-CsChrimson; dsx-DBD/+ |
| S9a | *D. melanogaster* | w; 15A01-p65.AD/+; 71G01-DBD/UAS-CD8::GFP |
| S9a | *D. yakuba* | w; 71G01-p65.AD/+; 15A01-DBD/UAS-CD8::GFP |
| S9b | *D. melanogaster* | w; dsx-Gal4/UAS-CD8::GFP |
| S9b | *D. yakuba* | w; dsx-Gal4/UAS-CD8::GFP |
| S9c | *D. melanogaster* | w; fru-Gal4/UAS-CD8::GFP |
| S9c | *D. yakuba* | w; fru-Gal4/UAS-CD8::GFP |
| S9c | *D. melanogaster* | LexAop-SPA-T2A-SPA/+; fru-LexA/+ |
| S9c | *D. yakuba* | UAS-SPA-GFP/+; fru-Gal4/+ |

**Table S2.**
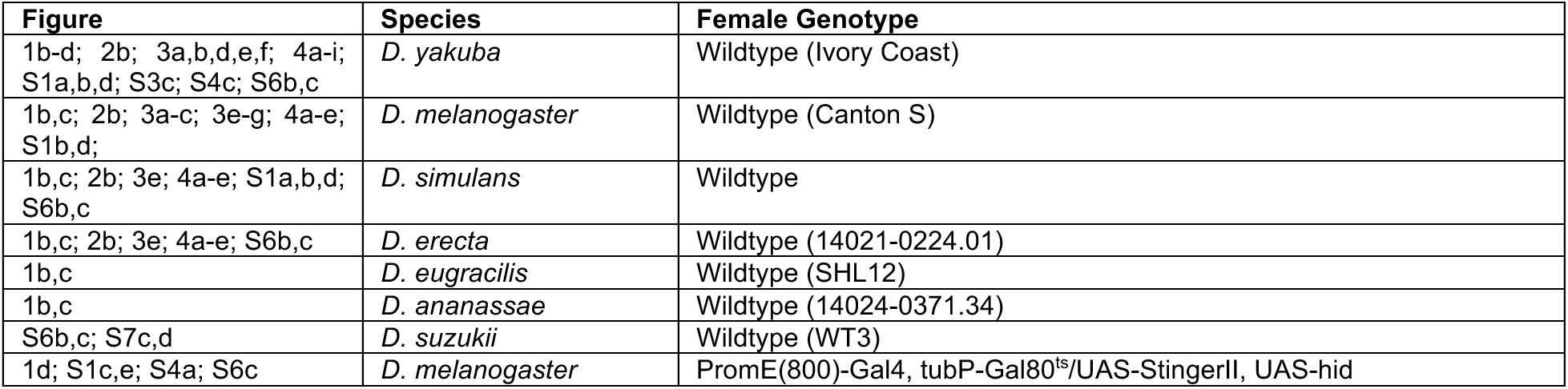
Female experimental genotypes by figure.

## Acknowledgements

We thank D. Stern for sharing several *D. yakuba* stocks; T. Hart and D. Kronauer for ie1 promoter DNA; M. Beye for hyperactive piggyBac transposase expression construct; B. Matthews and L. Seeholzer for technical advice; we thank D. Stern, B. Datta, B. Noro, J. Ouadah, A. Paul, T. Hindmarsh Sten, P. Brand, J. Rhee, A. Ryba and all the members of the Ruta laboratory for valuable discussion and comments on the manuscript. This work was supported by a NIH NINDS grant (5R35NS111611), the Simons Foundation Collaboration for the Global Brain and a Helen Hay Whitney Foundation Fellowship and NIH NIGMS grant (K99GM141319) for R.T.C. V.R. is a Howard Hughes Medical Institute Investigator.

## Author contributions

R.T.C. and V.R. conceived of the project. R.T.C., I.M., G.T.K., M.L.C., conducted and analyzed experiments. R.T.C., I.M., and Y.D. designed transgenesis strategies and built neurogenetic reagents. R.T.C. and V.R. wrote the manuscript. All authors provided critical feedback and helped shape the research, analysis and manuscript.

## Competing Interests

Authors declare that they have no competing interests.

## Data and materials availability

All data underlying this study are available upon request from the corresponding author.

## Notes

### Competing Interest Statement

The authors have declared no competing interest.

